# Hierarchical distribution matching enables comprehensive characterization of common and condition-specific cell niches in spatial omics data

**DOI:** 10.64898/2026.10.12.758505

**Authors:** Yuyao Liu, Jiashun Xiao, Xiaoheng Ma, Xiaomeng Wan, Peiqi Jiang, Zhiwei Wang, Yuheng Chen, Can Yang

## Abstract

Deciphering cell niches in complex tissues is essential for understanding tissue structure and disease. Recent advances in spatial omics have enabled subcellular resolution and accurate cell identity mapping. However, robust delineation of cell niches and disease-associated spatial patterns remains difficult. We introduce Harmonics, a novel computational framework that systematically identifies both common and condition-specific cell niches from spatial omics data through hierarchical distribution matching. Harmonics also includes a suite of downstream modules that facilitate comprehensive niche characterization. We demonstrate its scalability, accuracy and generalizability across datasets spanning diverse species, tissues, diseases, spatial modalities, and technological platforms. For condition-agnostic datasets, Harmonics outperforms baseline methods in both accuracy and robustness, and further demonstrates the capability to resolve niche structures at finer granularity. Across diverse diseases including pulmonary fibrosis, triple-negative breast cancer, and colorectal cancer, Harmonics enables precise identification of condition-specific niches and reveals disease-associated dynamics, subtype-specific spatial patterns, and structured immune architectures. We envision Harmonics as a practical and versatile tool for spatial niche analysis that can be applied across a wide range of biological contexts and seamlessly integrated with existing spatial omics workflows.

## Introduction

Cell niches (CNs) refer to microenvironments that typically exhibit unique cellular compositions and distinct interaction patterns, often associated with specific biological functions and regulatory programs^1–4^. Characterizing CNs enables the segmentation of tissue sections into functionally coherent regions, thereby focusing analysis on regions of interest (ROIs) while minimizing irrelevant signals and reducing computational burden. In case-control settings, disease tissue sections typically harbor both common niches shared with healthy samples that reveal normal tissue architectures, and condition-specific niches (CSNs) that more directly reflect pathogenic processes^5^. These CSNs warrant particular attention, as they may reveal therapeutic targets or prognostic markers^6^. Thus, a comprehensive understanding of both CNs and CSNs is crucial for decoding tissue function and disease mechanisms.

Spatial omics technologies, including MERFISH^7^, Xenium^8^, and Visium HD^9^ for transcriptomics, as well as IMC^10^, CODEX^11^, and MIBI-TOF^12^ for proteomics, have revolutionized tissue biology by preserving spatial context while capturing diverse biological features at single-cell or even subcellular resolution^13,14^. These technological advances have greatly expanded interest in CN and CSN discovery, creating an urgent need for computational methods capable of dealing with both settings. While existing methods for detecting CNs are available, they face notable limitations that hinder their practical utility. Several approaches^15–19^ first derive cell embeddings from profiled biological features, followed by clustering algorithms such as Leiden^20^ or mclust^21^ to define niches. The effectiveness and robustness of these methods heavily depend on the quality of the cell representations and the choice of clustering algorithm. However, due to substantial variability in data quality, resolution, and scale across different spatial omics platforms and modalities, such methods often lack generalizability, as producing reliable cell representations across diverse technologies remains inherently challenging. Moreover, most existing methods are not well-suited for characterizing CSNs under case-control settings. Some methods do not allow to incorporate condition-specific information for niche detection, leading to limited sensitivity to condition-associated signals^15–18^. Others oversimplify the task by detecting only a single CSN, overlooking the complex and heterogeneous nature of disease contexts^22,23^. Furthermore, certain methods cannot explicitly relate identified niches with specific conditions, thereby reducing biological interpretability^24^.

To overcome the aforementioned limitations and provide a reliable tool for delineating both CNs and CSNs, we propose Harmonics, a method that utilizes **H**ier**AR**chical distribution **M**atching for c**O**mprehensive characterization of common and condition-specific cell **NIC**hes in **S**patial omics data. Harmonics assumes that a cell’s neighborhood should exhibit a cell type distribution similar to that of its assigned niche if the assignment is biologically meaningful; otherwise, the cell is likely engaged in distinct biological processes and should not be grouped into that niche. Based on this assumption, Harmonics requires only the pre-defined cell type and spatial location of each cell as input. Cells and niches are both represented by cell type distributions, derived from their local neighborhoods and assigned cell members, respectively. This representation scheme allows Harmonics to be robustly applied to datasets across a variety of spatial omics platforms, unlike methods^15–19^ that rely on cell embeddings derived from noisy biological features. Building on this representation, we introduce hierarchical distribution matching (HDM) as the core algorithm of Harmonics, equipping it with the ability to characterize both CNs and CSNs in a unified framework. HDM alternates between identifying the optimal niche configuration at a fixed number of niches and merging the most similar niches, thereby generating a hierarchy of niche configurations across varying niche counts. Subsequently, Harmonics can automatically determine the optimal number of niches. Under the case-control setting, Harmonics begins by identifying basic cell niches (BCNs) from the control group. It then uses these BCNs as references and applies HDM to the case group to detect novel CSNs. Built on this computationally efficient framework, Harmonics is robust and readily scalable to datasets with millions of cells. It also includes comprehensive downstream modules for niche characterization to enhance its practical utility.

We demonstrate the superiority of Harmonics across a wide range of datasets encompassing different species, tissues, diseases, technologies, and omics modalities. Benchmarking on three datasets with expert-annotated niches shows that Harmonics outperforms baseline methods in both accuracy and robustness. We further apply Harmonics to a mouse brain hemisphere dataset with over two million cells, demonstrating its scalability and capacity to resolve CNs at fine granularity. In pulmonary fibrosis (PF), Harmonics uncovers disease-associated cell differentiation trajectories and captures dynamic alterations within a CSN correlated with disease progression. In triple-negative breast cancer (TNBC), it identifies a distinct CSN associated with patient prognosis. Finally, in two colorectal cancer (CRC) datasets from different modalities, Harmonics discovers shared immune niches and reveals spatial functional divergence among cells. We envision Harmonics as an accurate, robust, efficient, interpretable and generalizable tool that will serve as a go-to solution for niche-related research in spatial omics data.

## Results

### Overview of Harmonics

Harmonics is a unified framework designed to characterize both CNs and CSNs. It starts by constructing spatial relationships among all cells across all slices using Delaunay triangulation based on their spatial coordinates (Fig. 1a). This results in a neighborhood graph for each slice, from which a tailored cell neighborhood is defined for each cell (Methods). The cell type distribution within each neighborhood is then estimated and further used as the representation of the corresponding cell.

**Fig. 1.**
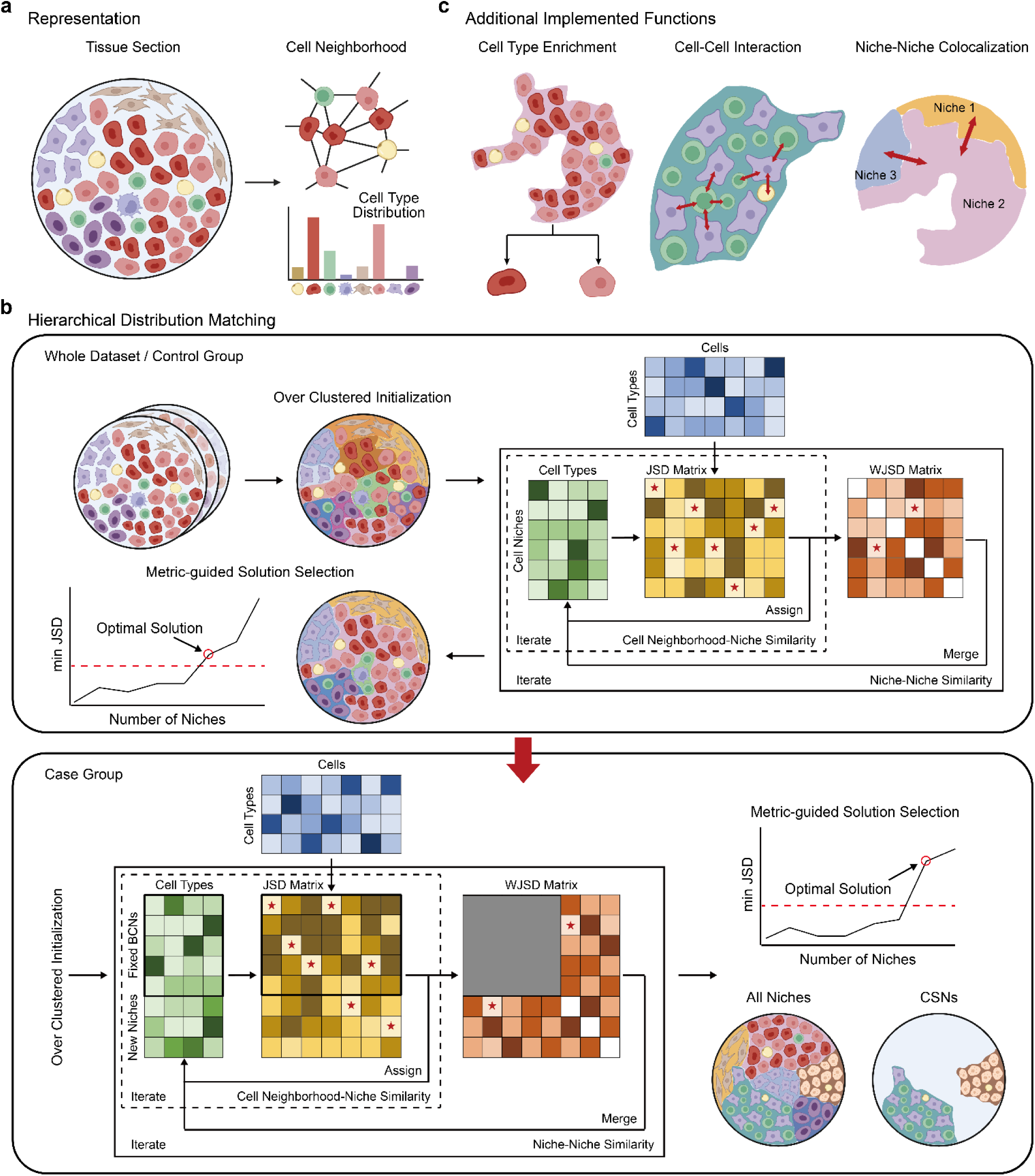
The overview of Harmonics. **a,** Harmonics constructs spatial relationships between cells within each slice based on their spatial coordinates using Delaunay triangulation, and further defines a cell neighborhood for each cell accordingly. The cell type distribution within each neighborhood is used as the representation of the corresponding cell. **b,** for condition-agnostic datasets, Harmonics identifies CNs by first over-clustering cells based on their representations, then applying its core algorithm HDM, which alternates between two steps: (i) iteratively reassigning each cell to the CN whose cell type distribution is most similar to that of its neighborhood (measured by JSD) and updating the cell type distribution of CNs until convergence; and (ii) merging the most similar CN pairs based on WJSD. The optimal CN number is selected using the *minJSD* score, ensuring niche separability without over-merging. In case-control settings, Harmonics first defines BCNs in the control group using the procedure described above, then applies HDM again to detect CSNs in the case group. During merging, BCNs remain fixed while CSNs are iteratively merged either with one another or into BCNs, with only the cell type distributions of CSNs being updated. The final CSN configuration is selected based on the *minJSD* score. **c,** Harmonics enhances niche characterization by incorporating downstream modules for cell type enrichment analysis, potential CCI identification, and NNC pattern detection.

For datasets that are condition-agnostic, Harmonics aims to identify CNs directly across the entire dataset (top panel of Fig. 1b). In this setting, Harmonics begins by over-clustering all cells based on their representations into a user-defined number of initial CNs, which is intentionally set to exceed the expected number of final CNs. Each CN is then represented by the cell type distribution of its assigned cells. Harmonics then applies HDM, its core algorithm, to produce a hierarchy of solutions by progressively merging similar CNs to obtain fewer representative CNs. The HDM process involves two alternating steps. First, Harmonics calculates the Jensen-Shannon divergence (JSD)^25,26^ between the cell type distribution of each cell’s neighborhood and that of each CN. Each cell is reassigned to the CN with the most similar distribution, and CN representations are updated accordingly. Harmonics repeats this process until convergence, resulting in a finalized cell-to-niche assignment for the current number of CNs. Second, Harmonics computes the weighted JSD (WJSD) between each pair of CNs, which accounts for both their size and similarity, and merges the most similar pair (Methods). These two steps are iterated, reducing the number of CNs until reaching a user-defined lower bound. Subsequently, Harmonics evaluates each candidate niche number using the *minJSD* score, which is defined as the smallest pairwise JSD between any two CNs in the corresponding solution. The final number of CNs is then selected as the point where *minJSD* first exceeds a predefined threshold. This reflects sufficient separability among all niche pairs without over-merging, and thus marking the optimal cell-to-niche assignment (Methods).

In case-control settings, where the samples are reasonably grouped into control and case cohorts, Harmonics aims to simultaneously identifies CSNs unique to the case group and BCNs that are shared between both groups (bottom panel of Fig. 1b). It first defines BCNs within the control group using the same HDM-based procedure described earlier. These BCNs are then used as fixed references during CSN detection in the case group. Harmonics performs over-clustering on the case group to initialize an additional set of candidate CSNs and re-applies the HDM procedure to obtain niche assignments across different CSN numbers. During the merging process, only CSNs are allowed to merge, either with one another or into BCNs, while the BCNs remain fixed and only the representations of the CSNs are updated (Methods). After all CSNs have been merged into BCNs, Harmonics uses the *minJSD* score to determine the optimal number of CSNs and generate the final cell-to-niche assignment. With its innovative algorithm design, Harmonics is computationally efficient for identifying both CNs and CSNs, and it readily scales to millions of cells in spatial omics data (Methods).

To achieve a comprehensive characterization of cell niches, Harmonics not only focuses on niche identification but also offers downstream modules for cell type enrichment analysis, identification of potential cell-cell interactions (CCIs), and detection of niche-niche co-localization (NNC) patterns (Fig. 1c). These modules quantify cell type enrichment within niches, the enrichment of neighboring cell type pairs, and the spatial co-occurrence of niche types (Methods), respectively, thereby enhancing the interpretability and biological characterization of the discovered niches.

### Harmonics outperforms existing methods in accurate and robust cell niche identification across brain datasets

We benchmarked Harmonics using several condition-agnostic spatially resolved transcriptomics (SRT) datasets generated by different technologies. To start with, we used a V1 cortex slice generated by STARmap (STARmap V1C)^27^, which was annotated by experts into seven distinct CNs. Prior to analysis, we removed smooth muscle cells, as they are typically associated with the pial surface but were rarely present in this dataset and had been roughly merged into layer 1 in the original annotation. We compared Harmonics against seven baseline methods that are also condition-agnostic (Methods). Among all methods, Harmonics best captured the spatial structure of all seven annotated CNs. For instance, only Harmonics and scNiche^18^ were able to identify layer 1. However, scNiche was unable to distinguish the corpus callosum from the hippocampus, erroneously merging them into a single CN, while Harmonics correctly resolved them as separate niches (Fig. 2a). To systematically evaluate performance, we ran each method 20 times with different random seeds and evaluated the results based on four commonly used metrics, namely Adjusted Rand Index (ARI), Adjusted Mutual Information (AMI), macro F1-score (mF1), and weighted F1-score (wF1) (Methods). Harmonics consistently outperformed all baseline methods across all metrics (Fig. 2b, Supplementary Fig. 1a).

**Fig. 2.**
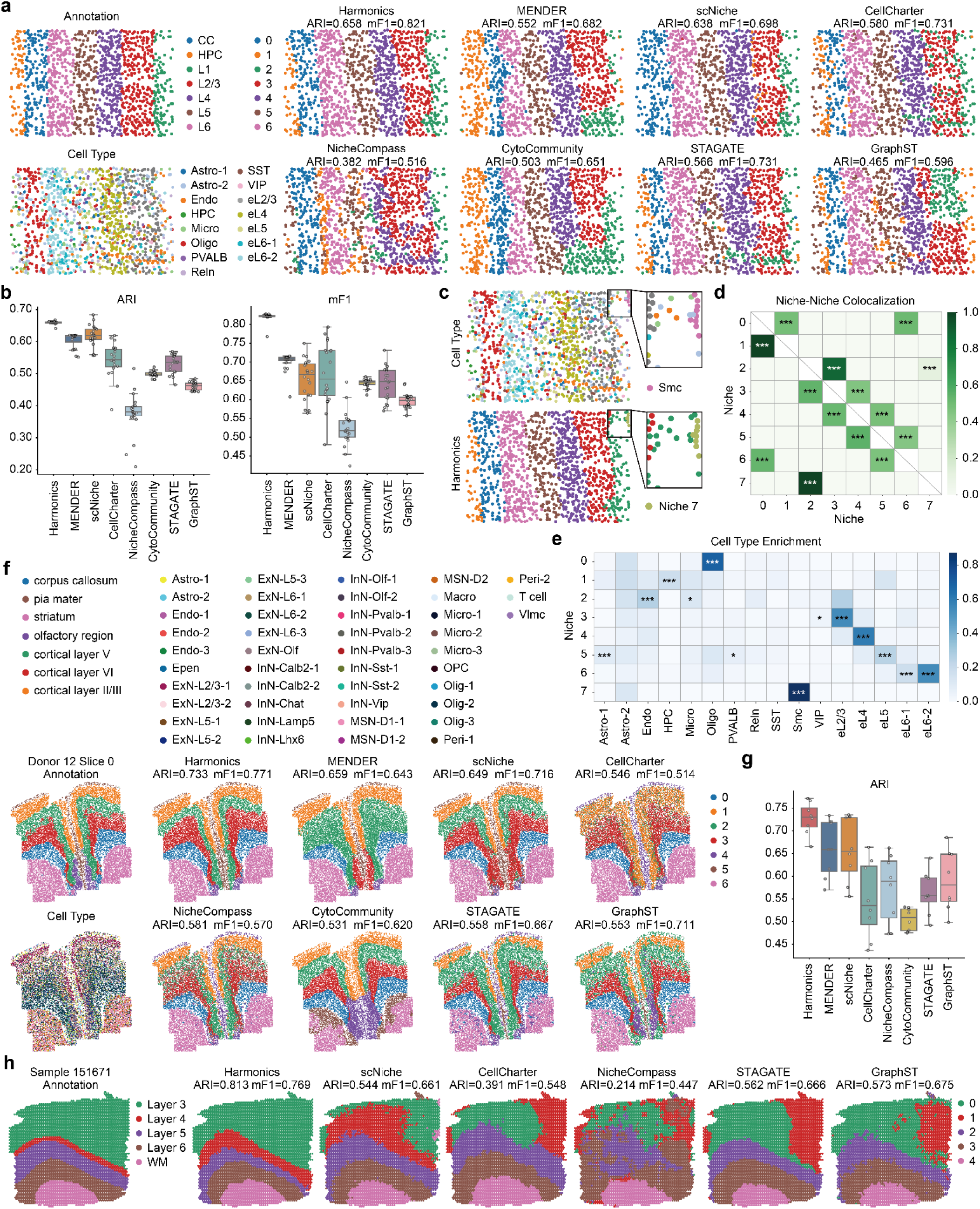
Harmonics exhibits accurate and robust CN identification across brain datasets. **a,** the single slice of the STARmap V1C dataset, colored by niche annotation, cell type, and the matched CN assignments by Harmonics and baseline methods in a representative run. Corresponding ARI and mF1 score are displayed. CC, corpus callosum; HPC, hippocampus; L1 to L6, the six neocortical layers. **b,** comparison of ARI and mF1 scores over 20 independent runs on the STARmap V1C dataset, showing Harmonics outperforms all baseline methods. Each dot represents the outcome of one run. **c,** CN detection results produced by Harmonics with smooth muscle cells ("Smc") included and the optimal CN counts automatically determined by Harmonics. A zoomed-in view highlights the region including smooth muscle cells, with slices colored by cell type (top) and Harmonics results (bottom). **d,** NNC patterns identified by Harmonics on the STARmap V1C dataset, accurately reflecting spatial organization of distinct niches. Rows represent source niches and columns represent target niches. A significant value indicates that the corresponding target niche is significantly enriched in the spatial neighborhood of the corresponding source niche. **e,** cell type enrichment results for STARmap V1C dataset, revealing biologically coherent enrichment patterns across niches. **f,** a representative slice from the 24-week-old group of the MERFISH FC&S dataset, colored by niche annotation, cell type, and the CN assignments by Harmonics and baseline methods, along with their respective ARI and mF1 scores. **g,** comparison of ARI scores, where Harmonics outperforms all baseline methods. Each dot represents results of an individual slice from the 24-week-old group of the MERFISH FC&S dataset. **h,** a representative slice from the Visium DLPFC dataset, colored by niche annotation and CN assignments from Harmonics and baseline methods, with corresponding ARI and mF1 scores. For all box plots, the center line indicates the median; box limits represent the upper and lower quartiles; whiskers extend to the largest and smallest values within 1.5 times the interquartile range. *, adjusted *p*-value < 0.05; **, adjusted *p*-value < 0.01; ***, adjusted *p*-value < 0.001.

The key to the success of Harmonics stems from both its hierarchical framework and its JSD-based cell-to-niche assignment strategy. To see this, we compared Harmonics to two alternative strategies. The first involved directly clustering the cell representations derived by Harmonics into seven clusters using k-means. The second retained the hierarchical framework of Harmonics but replaced the first step of HDM with k-means clustering at each candidate number of CNs. Harmonics outperformed both alternatives (Supplementary Fig. 1a). Interestingly, even the second alternative outperformed all baseline methods and the first alternative, emphasizing the advantage of the hierarchical design.

We evaluated the robustness of each method by measuring the consistency between pairs of repeated runs (Methods). Harmonics demonstrated higher robustness compared to all baseline methods (Supplementary Fig. 1b). Interestingly, MENDER^28^ and CytoCommunity^24^, which ranked just below Harmonics in robustness, were the only baselines that exclusively used cell type information for CN identification. This highlights the advantage of cell type-based inputs in mitigating noise from molecular profiling and achieving more stable CN detection.

Subsequently, we included smooth muscle cells in the analysis and allowed Harmonics to automatically determine the number of niches. Harmonics identified eight CNs, among which the original seven annotated CNs were preserved (Supplementary Fig. 1c). In addition, the spatially clustered smooth muscle cells were identified as a distinct CN (Fig. 2c). Based on this result, we evaluated the performance of the capability of Harmonics in NNC pattern detection and cell type enrichment analysis through its implemented modules. The NNC pattern detection module accurately reflected the spatial organization of the V1C region. Specifically, all CNs were only enriched for connections with their immediate neighboring CNs, capturing the layered cortical structure (Fig. 2d). Moreover, Harmonics effectively revealed the dominant cell types within each CN using the cell type enrichment module. For instance, excitatory neurons specific to each neocortical layer were found to be enriched only in the corresponding CN (Fig. 2e).

In a MERFISH dataset of mouse frontal cortex and striatum (MERFISH FC&S)^1^, comprising three groups corresponding to different developmental stages (4-week-old, 24-week-old, and 90-week-old), Harmonics again outperformed all baseline methods. Taking a representative slice from the 24-week-old group as an example, Harmonics accurately identified seven CNs that were highly consistent with expert annotations. Among all methods, only Harmonics and STAGATE^15^ were able to fully recover both layer 5 and layer 6. However, Harmonics showed superior performance in identifying the pia mater and striatum, capturing these regions as complete and spatially coherent CNs, in contrast to the fragmented results produced by STAGATE (Fig. 2f). Quantitative evaluation confirmed the consistently superior performance of Harmonics over all baseline methods across the three groups of data (Fig. 2g, Supplementary Fig. 1d). Furthermore, we evaluated JSD against other commonly used metrics for measuring distributional similarity, using the STARmap V1C and MERFISH FC&S datasets (Methods), and found that JSD offers a better balance between accuracy and robustness in niche identification (Supplementary Fig. 2).

We further evaluated the ability of Harmonics to generalize to low-resolution SRT data using the Visium human dorsolateral prefrontal cortex dataset (Visium DLPFC)^29^, which comprises three groups of samples. We applied our previously developed method, STitch3D^30^, to perform cell type deconvolution and used the resulting cell type compositions of spots as input for Harmonics, instead of cell type labels, due to the resolution limitation (Methods). We then compared the performance of Harmonics against baseline methods suitable for low-resolution data. Harmonics outperformed all baseline methods for samples in groups 1 and 2, and achieved comparable performance to STAGATE and GraphST^16^ for samples in group 3, while still surpassing the other baselines (Supplementary Fig. 3). Notably, Harmonics more accurately captured the laminar architecture of the cortex. For instance, only Harmonics successfully identified layer 4 in samples from group 2, whereas other methods either missed this layer or produced patterns that deviated substantially from the annotated morphology (Fig. 2h).

Through a comprehensive set of benchmarks, we demonstrated that Harmonics achieves superior performance in both CN characterization and robustness across a wide range of datasets generated by diverse spatial omics technologies, underscoring its broad applicability.

### Harmonics scales to complex brain-wide atlases and enables discovery of fine-grained niche structures

To evaluate both the scalability and the capacity of Harmonics to identify fine-grained CNs, we applied Harmonics to a large-scale MERFISH dataset comprising 23 consecutive sagittal slices from a whole mouse brain hemisphere (MERFISH WMB)^31^. The dataset contains over two million cells annotated into more than 300 distinct cell types. Several graph neural network (GNN)-based methods, including CytoCommunity, STAGATE, and GraphST, are not applicable to this dataset with millions of cells, suggesting the scalability issue of graph-based methods for analyzing large-scale SRT datasets.

Using its metric-guided CN number selection strategy, Harmonics identified 61 CNs on this dataset (Supplementary Fig. 4). In contrast, scNiche and CellCharter^17^, which also incorporate an automatic CN number determination procedure, identified only 11 and 4 CNs, respectively. This indicates that they may not be suitable to capture the fine-grained structural complexity inherent in this large-scale dataset. MENDER requires a user-specified target CN number and employs resolution search to determine the final result. As it lacks an internal strategy for selecting the optimal CN count, we set the target to 61 to ensure a fair comparison. Nonetheless, MENDER stopped at 36 CNs and was still unable to match the granularity achieved by Harmonics (Fig. 3a).

**Fig. 3.**
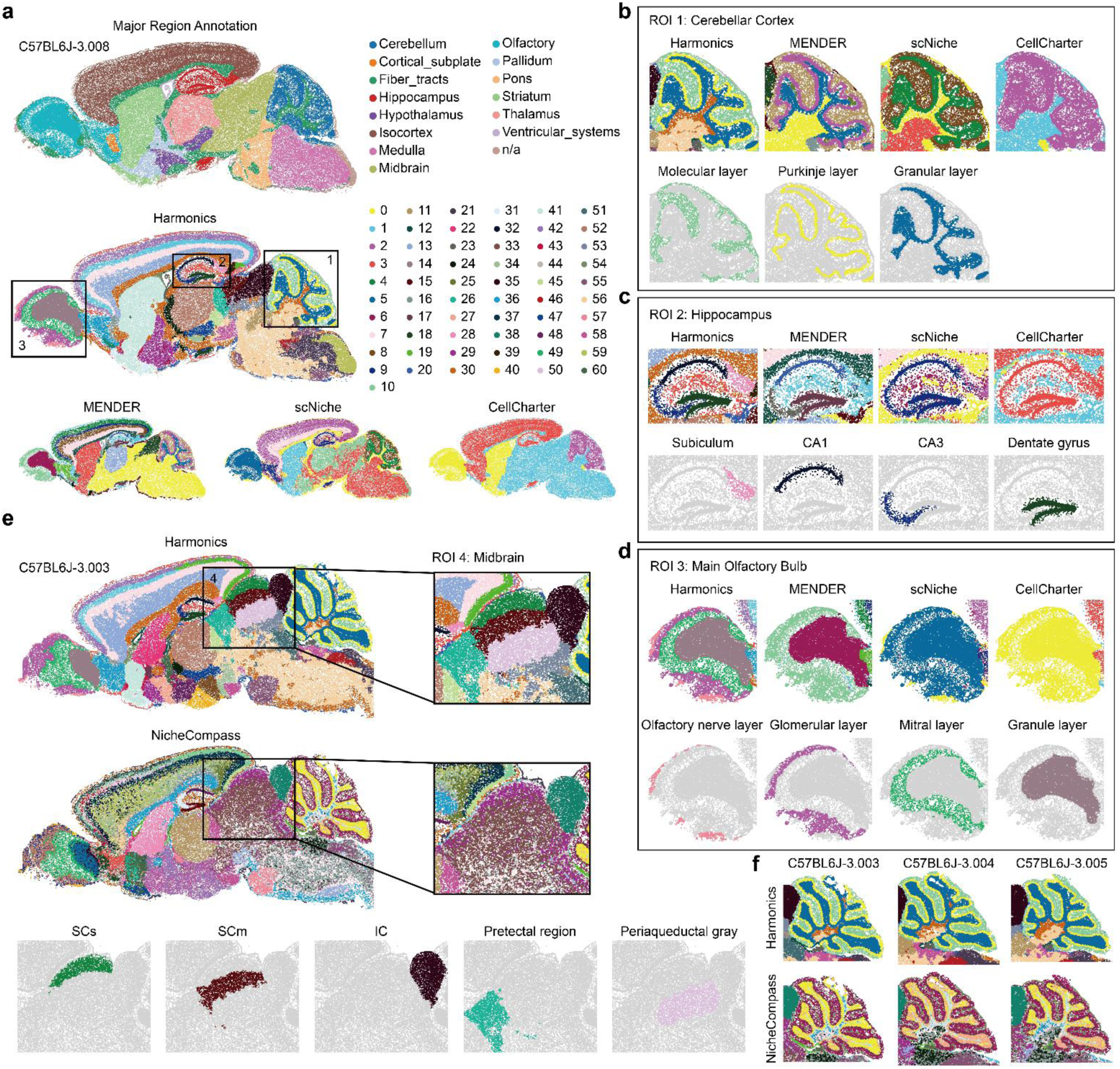
Harmonics demonstrates the ability to identify fine-grained CNs in whole mouse brain hemispheres. **a,** comparison on a representative slice from the MERFISH WMB dataset between Harmonics and other methods that support automatic CN number selection (CellCharter and scNiche), as well as MENDER, which determines the final results through a resolution search strategy guided by a pre-specified number of target niches. Harmonics identifies 61 CNs, while scNiche and CellCharter only identify 11 and 4 CNs, respectively; MENDER stops at 36 CNs. Three ROIs are highlighted. **b**-**d,** comparison of results in three representative brain structures, including cerebellar cortex (**b**), hippocampus (**c**), and main olfactory bulb (**d**). For each region, the top panel shows a comparison between Harmonics and baseline methods, while the bottom panel highlights the major CNs identified by Harmonics within the corresponding structure. **e,** comparison between Harmonics and NicheCompass in the midbrain region on a representative slice. Harmonics identifies five biologically meaningful CNs, whereas NicheCompass does not resolve such fine-grained structures. SCs, superior colliculus, sensory related; SCm, superior colliculus, motor related; IC, inferior colliculus. **f,** comparison between Harmonics and NicheCompass in the cerebellar cortex across three consecutive slices, where Harmonics yields more consistent CN definitions across slices than NicheCompass.

The dataset includes coarse annotations for 15 major brain regions (Fig. 3a). We first examined the distribution of each CN identified by Harmonics across these annotated brain regions, with one unnamed region excluded from the analysis. The majority of CNs were predominantly localized within a single major brain region, supporting the spatial specificity and biological interpretability of the CN definitions (Supplementary Fig. 5a). We then focused on several anatomically well-defined regions of interest (ROIs) and compared Harmonics with baseline methods. The CNs identified by Harmonics within these ROIs were validated based on representative enriched cell types identified through the cell type enrichment analysis (Supplementary Figs. 5 and 6), as well as anatomical annotations from the Allen Brain Atlas^32^. In the cerebellar cortex, Harmonics accurately delineated the three-layered structure comprising the molecular layer, Purkinje layer, and granular layer. MENDER also recovered this organization, while scNiche and CellCharter coarsely grouped this region into only two and one CNs, respectively (Fig. 3b, Supplementary Fig. 5b and c). In the hippocampus, Harmonics distinguished between CA1, CA3, and dentate gyrus, whereas scNiche and CellCharter merged these subregions into a single CN. Although MENDER was also able to identify these three subregions, it did not capture the subiculum, which Harmonics successfully identified (Fig. 3c, Supplementary Fig. 5d and e). In the main olfactory bulb, Harmonics further subdivided this region into four CNs, namely the olfactory nerve layer, glomerular layer, mitral layer, and granule layer, whereas all three baseline methods identified at most two CNs in this area (Fig. 3d, Supplementary Fig. 5f and g). In addition, Harmonics identified three distinct CNs in the ventricular system, enriched for ependymal cells, non-neuronal choroid plexus cells, and a specific kind of inhibitory immature neurons^31^, respectively (Supplementary Fig. 6a to c).

We further compared Harmonics with NicheCompass^19^, which defines CNs by directly clustering its learned representations without automatic selection of the optimal CN number. We clustered its output into 61 categories, matching the number of CNs found by Harmonics (Methods). Although NicheCompass generally identified finer-grained CNs than other baseline methods, it was unable to resolve important spatial structures in specific regions. In the midbrain, Harmonics accurately identified five distinct CNs corresponding to the sensory-related superior colliculus, motor-related superior colliculus, inferior colliculus, pretectal region, and periaqueductal gray. In contrast, NicheCompass captured only the inferior colliculus, while other regions appeared as mixed categories lacking clear spatial boundaries. Similar observations were made with other baselines (Fig. 3e, Supplementary Fig. 6d to f). Harmonics also revealed well-separated CNs with clear boundaries in the hypothalamus, while NicheCompass did not resolve the same level of niche diversity (Supplementary Figs. 6g). Additionally, Harmonics exhibited greater consistency in CN definitions across adjacent slices of the same brain region, a desirable quality that NicheCompass did not achieve (Fig. 3f). These results suggest that the cell embeddings learned by NicheCompass lack sufficient resolution to support the identification of fine-grained CNs comparable to Harmonics.

Harmonics also incorporates a label transfer module to improve its efficiency on large-scale datasets with high inter-slice homogeneity (Methods). Specifically, we selected one out of every four slices to define CNs and directly assigned cells from the remaining slices to these CNs via label transfer. We further evaluated the consistency between the label transfer-based assignments and those obtained directly from the full dataset (Methods). The high agreement (median FMI of 0.93 and mean FMI of 0.92) underscores the robustness of Harmonics (Supplementary Figs. 6h).

### Harmonics uncovers meaningful CSNs and disease-associated dynamics in pulmonary fibrosis

To further evaluate the ability of Harmonics to identify CSNs in the case-control setting, we applied it to a Xenium dataset, which includes distal lung sections from both healthy donors and patients with PF (Xenium PF)^2^. Each slice in this dataset is annotated with a visually estimated pathology percentage by pathologists to indicate the extent of tissue remodeling. We designated the healthy lung slices as the control group and the PF lung slices as the case group. Harmonics first identified four distinct BCNs in the control group. These BCN definitions were then fixed and used as a reference for the case group, where Harmonics additionally discovered four CSNs (Supplementary Fig. 7a). Comparing niches between groups revealed that each BCN from the control group corresponded closely to its counterpart in the case group, with minimal similarity to other BCNs or any CSNs (Fig. 4a, Methods). This result indicates that although BCN definitions were not updated during CSN detection, Harmonics was still able to achieve reasonable niche partitioning for regions in the case group that resemble those in the control group, thereby supporting the validity of its underlying assumptions (Supplementary Fig. 7b).

**Fig. 4.**
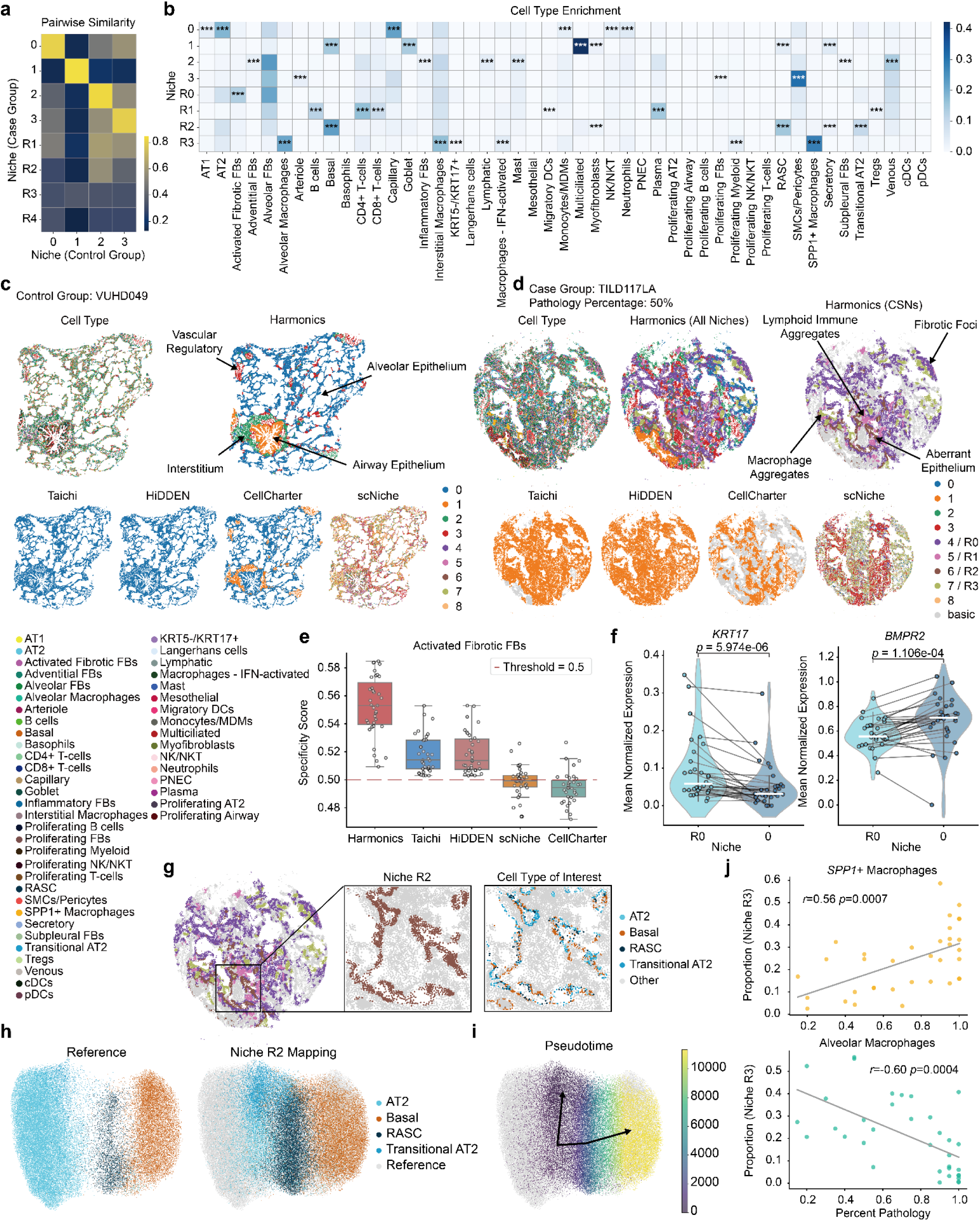
Harmonics reveals disease-associated dynamics of CSNs in pulmonary fibrosis. **a,** similarity between each BCN from the control group and each BCN/CSN in the case group, quantified by 1-JSD. Higher scores indicate greater similarity in cell type distributions between the two niches. **b,** cell type enrichment analysis of the case group. CSNs defined by Harmonics are labeled with the prefix "R". *, adjusted *p*-value < 0.05; **, adjusted *p*-value < 0.01; ***, adjusted *p*-value < 0.001. **c,** a representative slice from the control group, colored by cell type and niche assignments from Harmonics and baseline methods. Representative regions corresponding to the four BCNs identified by Harmonics are highlighted. **d,** a representative slice from the case group, shown with cell type annotations, all niches identified by Harmonics, and CSNs defined by both Harmonics and baseline methods, where "basic" denotes niches that are not recognized as CSNs. Representative regions corresponding to the four CSNs discovered by Harmonics are highlighted. **e,** comparison of the specificity of activated fibrotic FBs enrichment in CSNs across all methods. Each dot represents a single slice. For each method, the CSN with the highest median specificity across all slices is used for comparison. Dots below the red dashed line represent slices in which the CSN has a lower proportion of activated fibrotic FBs than the rest of the tissue, indicating poor specificity. For the box plots, the center line indicates the median; box limits represent the upper and lower quartiles; whiskers extend to the largest and smallest values within 1.5 times the interquartile range. **f,** two representative DEGs of AT2 cells between CSN-R0 and BCN-0 in the case group. Each dot represents a slice, and *p*-values are calculated using a two-sided Wilcoxon signed-rank test. **g,** a representative view of CSN-R2, showing the spatial distribution of four major epithelial cell types: AT2, basal, RASC, and transitional AT2. **h,** UMAP of AT2, basal, RASC, and transitional AT2 cells from BCN-0 and BCN-1 used as a reference (left), and the projection of the same four cell types from CSN-R2 onto the reference UMAP as query data (right). **i,** trajectory and pseudotime inference of the query data shown in (**h**), with cells colored by pseudotime. The black lines depict two differentiation trajectories inferred for AT2 cells. **j,** correlation between pathology percentage and the proportions of two major macrophage populations in CSN-R3. Each dot represents a slice. Trend lines are fitted using univariate linear regression, and Pearson’s *r* along with *p*-values from a two-sided permutation test are reported.

Cell type enrichment analysis allowed us to examine the biological properties of both the identified BCNs and CSNs (Fig. 4b). In the control group, we observed well-organized structural niches corresponding to distinct anatomical compartments of the lung (Fig. 4c, Supplementary Fig. 7c). BCN-0 was enriched for alveolar type 1 (AT1) and type 2 (AT2) cells, which are the main epithelial cell types lining the alveolar surface and are essential for gas exchange and surfactant production^33^. This niche also contained a small number of homeostatic immune cells, reflecting the immune surveillance in the alveoli. BCN-0 dominated the spatial landscape in the control group and represents the alveolar epithelium. BCN-1 was enriched for cell types such as multiciliated cells, basal cells, goblet cells, and respiratory airway secretory cells^34,35^ (RASCs), with clear anatomical localization along bronchial structures, representing the airway epithelium. BCN-2 featured multiple subtypes of fibroblasts (FBs), including alveolar FBs, adventitial FBs, and subpleural fibroblasts FBs, which together constituted the interstitial compartment. BCN-3 was enriched for pericytes and smooth muscle cells, representing the vascular regulatory region that supports vascular tone, capillary stabilization, and tissue homeostasis^36^.

The four CSNs identified by Harmonics in the case group also exhibited distinct biological features (Fig. 4d, Supplementary Fig. 8). CSN-R0 was specifically enriched for activated fibrotic fibroblasts and represented fibrotic foci, predominantly occupying the remodeled alveolar interstitium. CSN-R1 was enriched for multiple immune cell types, including B cells and several types of T cells, corresponding to lymphoid immune aggregates that are rarely present in healthy lungs. CSN-R2, defined as aberrant epithelium, was enriched for transitional AT2 cells, together with two airway epithelial cell types, basal cells and RASCs, which abnormally appeared in alveolar regions. CSN-R3 represented macrophage aggregates located within alveolar and airway lumens, a feature not observed in the healthy lungs.

We benchmarked Harmonics against Taichi^22^ and HiDDEN^23^, two methods specifically developed for detecting condition-relevant niches or cell populations. Additionally, we also included scNiche and CellCharter, which can automatically determine the number of spatial niches. For these two methods, we considered niches with higher proportions in the case group than in the control group as disease-relevant or condition-relevant (Methods). We also attempted to include CytoCommunity, which offers a supervised mode specially for case-control studies; however, it could not handle the full Xenium PF dataset due to the out-of-memory error. We found that only Harmonics, CellCharter, and HiDDEN showed significant correlations between overall CSN proportions and pathology percentage (Supplementary Fig. 9a, Methods). However, HiDDEN classified nearly the entire tissue area in diseased slices as CSN, with the lowest CSN proportion still around 80%, even in slices where the pathology percentage was below 20%. In addition, non-CSN cells defined by HiDDEN were dispersed rather than forming coherent structures, indicating poor spatial specificity (Supplementary Fig. 8). We further evaluated the ability of each method to enrich disease-associated cell types within CSNs, including activated fibrotic fibroblasts, transitional AT2 cells, and B cells (Methods). Harmonics consistently demonstrated the highest specificity (Fig. 4e, Supplementary Fig. 9b). Notably, CellCharter only identified two niches across slices and assigned most activated fibrotic fibroblasts and transitional AT2 cells to the niche more abundant in the control group, suggesting that it may not fully capture PF-associated structures. Moreover, baseline methods also struggled to delineate well-structured niches within the control group (Fig. 4c). In conclusion, Harmonics outperformed all baseline methods in the case-control setting by effectively identifying both BCNs and CSNs.

We next focused on individual CSNs to examine disease-associated processes. In CSN-R0 (fibrotic foci), we observed the presence of a subset of AT2 cells. Differential expression analysis (Methods) comparing these cells to their counterparts in BCN-0, representing alveolar epithelium, revealed that AT2 cells in CSN-R0 significantly upregulated *KRT17* and *TGFB2*, both associated with TGF-β pathway activation^37–39^. Conversely, they downregulated *SFTPC* and *BMPR2*, which are essential for mature AT2 identity and BMP signaling^39–41^, respectively (Fig. 4f, Supplementary Fig. 9c). These changes indicate that AT2 cells in fibrotic foci are under aberrant regulation by heightened TGF-β signaling and suppressed BMP signaling. Consistently, activated fibrotic FBs in CSN-R0 expressed markedly higher levels of *TGFB1* and *CTHRC1* compared to normal alveolar FBs^42,43^ (Supplementary Fig. 9d). Both are potent TGF-β pathway activators and are likely to promote partial epithelial-to-mesenchymal transition (EMT)-like changes in neighboring AT2 cells^44^ and drive phenotypic conversion of AT2 cells toward a transitional state (Supplementary Fig. 9c). Moreover, these FBs expressed elevated levels of *SFRP2*, *SFRP4*, and *WNT5A* (Supplementary Fig. 9d), which modulate Wnt signaling and have been shown to suppress BMP signaling^45–48^. These effects may limit the ability of AT2 cells to differentiate into AT1 cells^49^ and contribute to the pathological accumulation of transitional AT2 cells in fibrotic foci, where they fail to attain mature alveolar identity^50–52^. Supporting this mechanism, the CCI detection module in Harmonics revealed enriched spatial proximity of AT2 cells to both activated fibrotic FBs and transitional AT2 cells (Supplementary Fig. 10). This spatial enrichment further suggests that aberrant signaling between fibroblasts and epithelial cells may contribute to the altered AT2 cell states observed in fibrotic lungs.

In CSN-R2, which represents aberrant epithelium, we observed a spatial co-localization of alveolar epithelial cell types, such as AT2 and transitional AT2, with airway epithelial cell types, including basal cells and RASCs. While these cell types typically occupy distinct anatomical compartments in healthy lungs, they appeared highly intermixed in CSN-R2 (Fig. 4g). CCI detection revealed enriched spatial adjacency among all four cell types, indicating potential inter-lineage interactions (Supplementary Fig. 11). We found that AT2 cells in CSN-R2 showed reduced expression of canonical alveolar markers and increased expression of genes associated with airway epithelial programs compared to AT2 cells in BCN-0 (Supplementary Fig. 9e), suggesting a shift toward secretory-like or basal-like phenotypes^35,53^. To further investigate this hypothesis, we constructed a reference UMAP using AT2, transitional AT2, basal, and RASC cells from BCN-0 and BCN-1, which revealed clear separation between alveolar and airway cell types (Fig. 4h, Supplementary Fig. 12a). When projecting CSN-R2 cells onto this reference (Methods), we observed that AT2 cells in CSN-R2 shifted toward the basal and RASC clusters, consistent with a phenotypic transition toward airway-like states (Fig. 4h, Supplementary Fig. 12a and b). Trajectory inference and pseudotime analysis within CSN-R2 identified two major differentiation paths (Fig. 4i, Methods). One path extended from AT2 cells through RASCs to basal cells, suggesting a potential mechanism for bronchiolization of alveolar regions during fibrosis^35^. This trajectory also aligns with the general trend of increased basal cell proportion in CSN-R2 as disease severity increases (Supplementary Fig. 12e, Methods). The other path represented a transition from AT2 to transitional AT2. Given that CSN-R2 is primarily adjacent to fibrotic foci (Supplementary Fig. 12d), it is plausible that similar signaling abnormalities contribute to the observed transitions, including the accumulation of transitional AT2 cells. These differentiation pathways were further validated by combined analysis of BCN-0, BCN-1, and CSN-R2 (Supplementary Fig. 12c).

Furthermore, we observed that the relative abundance of *SPP1*+ macrophages and alveolar macrophages specifically within CSN-R3 (macrophage aggregates) was significantly associated with pathology percentage. As the severity of pathology increased, the proportion of *SPP1*+ macrophages in CSN-R3 tended to rise, while alveolar macrophages showed a decreasing trend. This shift may reflect a disease-driven replacement of homeostatic macrophages by fibrotic or proinflammatory subtypes^54^. Interestingly, the overall proportion of *SPP1*+ macrophages across the entire tissue section did not show a significant association with pathology percentage (Supplementary Fig. 12f). This highlights the advantage of CSN-based analysis in revealing disease-associated dynamics.

### Harmonics reveals subtype-specific spatial patterns and prognostic niche architecture in triple-negative breast cancer

We applied Harmonics to a proteomics dataset generated by the MIBI-TOF technology which consists of tissue sections from 40 patients with TNBC (MIBI-TOF TNBC)^12^ to evaluate its applicability beyond transcriptomic modalities. Although all samples are derived from the same cancer type, they exhibit notable heterogeneity. A subset of samples contains very few immune cells and is designated as the cold group, while the remaining samples are further classified into the compartmentalized group, where immune and tumor components are spatially separated, and the mixed group, where immune and tumor cells are intermingled^12^ (Methods). To identify CSNs, we treated the cold group as the control group and merged the compartmentalized and mixed groups into the case group without distinguishing between the two subtypes. Harmonics identified BCNs primarily composed of various combinations of tumor and non-immune cell types (Fig. 5a, Supplementary Fig. 13). In contrast, the CSNs represented distinct immune components (Fig. 5a and b, Supplementary Fig. 13b). Given that CSN-R4 is enriched for multiple immune cell types, including neutrophils, macrophages, CD8+ T cells, and regulatory T cells (Tregs), and concurrently contains a substantial proportion of Keratin+ tumor cells, the predominant tumor cell type in TNBC, it likely represents a tumor-immune interaction niche.

**Fig. 5.**
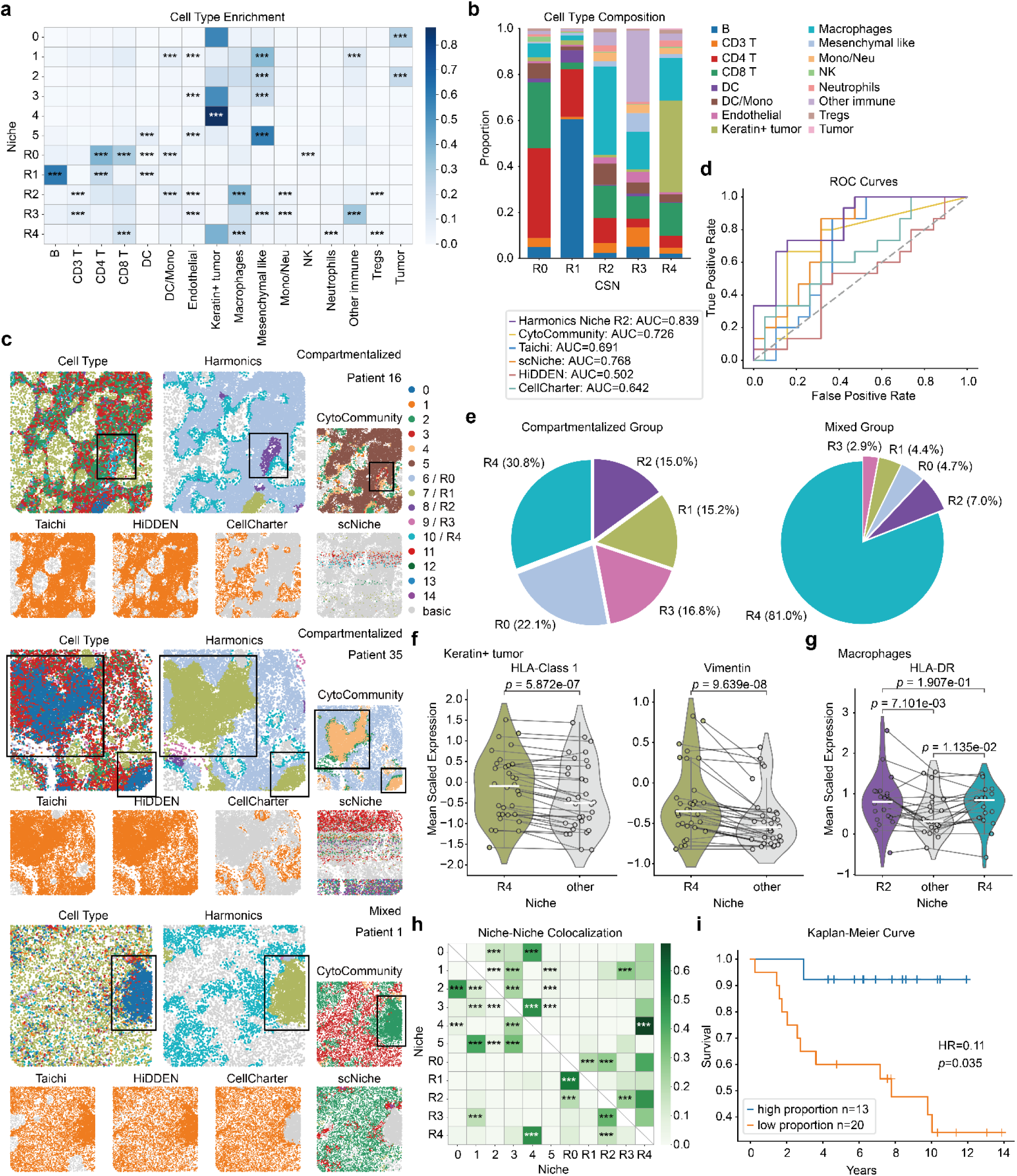
Harmonics depicts distinct immune-associated CSNs in triple-negative breast cancer. **a,** cell type enrichment analysis for the case group in the MIBI-TOF TNBC dataset. CSNs identified by Harmonics are labeled with the prefix "R". **b,** cell type compositions of different CSNs in the case group. **c,** representative slices from the compartmentalized and mixed groups, colored by cell types and CSNs identified by Harmonics and baseline methods, where "basic" denotes niches not recognized as CSNs. The boxed ROIs highlight regions where Harmonics captures CSNs and their spatial consistency across slices more effectively than CytoCommunity. **d,** comparison of different methods in distinguishing compartmentalized and mixed groups using niche proportion. The AUC is reported. For Harmonics, the proportion of CSN-R2 is used to classify disease subtypes; for other methods, the niche that can best distinguish between subtypes is selected. **e,** pie charts showing the composition of CSNs in the compartmentalized and mixed groups. The compartmentalized group exhibits relatively balanced proportions of five CSNs, whereas the mixed group is dominated by CSN-R4. **f,** differential expression of HLA-Class 1 and Vimentin in Keratin+ tumor cells between CSN-R4 and other niches. **g,** differential expression of HLA-DR in macrophages between CSN-R2/CSN-R4 and other niches. In panels **f** and **g**, each dot represents a slice, and *p*-values are computed using a two-sided Wilcoxon signed-rank test. **h,** NNC patterns identified by Harmonics in the MIBI-TOF TNBC dataset. Rows represent source niches and columns represent target niches. A significant value indicates that the target niche is significantly enriched in the spatial neighborhood of the source niche. **i,** Kaplan-Meier survival curves for patients stratified by high and low intra-slice proportions of CSN-R2. The hazard ratio (HR) is computed using Cox regression and the associated *p*-value is obtained from a two-sided test. The threshold for stratification in this plot is 0.03. *, adjusted *p*-value < 0.05; **, adjusted *p*-value < 0.01; ***, adjusted *p*-value < 0.001.

We compared the CSNs identified by Harmonics with those obtained from baseline methods (Fig. 5c, Supplementary Figs. 14 and 15). In this dataset, scNiche did not produce well-structured results, while CellCharter only identified two niches and predominantly assigned immune cells to a niche that was more abundant in the control group, indicating that condition-agnostic methods have limited ability to delineate CSNs. Taichi and HiDDEN demonstrated relatively better performance in recognizing immune components as condition-specific. However, both methods are inherently limited to binary classification, restricting CSN identification to a single immune-associated niche and thus lacking the ability to resolve distinct immune components. In contrast, Harmonics and CytoCommunity demonstrated superior capacity to identify immune-enriched CSNs. For example, in the top panel of Fig. 5c, both methods successfully distinguished a T cell-enriched niche (CSN-R0 in Harmonics and CN5 in CytoCommunity). However, compared to CytoCommunity, Harmonics additionally identified a macrophage-dominant region (CSN-R2), which CytoCommunity did not capture. Moreover, the CSNs defined by Harmonics exhibited better inter-slice consistency. For instance, in the middle and bottom panels of Fig. 5c, Harmonics consistently recognized a B cell-enriched region as CSN-R1 across two different slices, whereas CytoCommunity assigned them to inconsistent niche identities. We also observed that CSN-R4, identified by Harmonics, localized to tumor boundaries in the compartmentalized group but formed irregular regions in the mixed group (Supplementary Figs. 14 and 15).

Furthermore, we evaluated the ability of each method to distinguish between the compartmentalized and mixed subtypes using on niche proportions (Methods). Harmonics achieved higher area under the ROC curve (AUC) scores using both CSN-R2 and CSN-R0 compared to the top-performing niches from baseline methods (Fig. 5d, Supplementary Fig. 16a and b). In addition, the composition of CSNs differed markedly between the two subtypes. In the compartmentalized group, the proportions of different CSNs were relatively balanced, whereas in the mixed group, CSN-R4 was overwhelmingly dominant (Fig. 5e). These results further highlight the superior capability of Harmonics in capturing subtype-specific spatial patterns.

We found that the same cell type differs in protein expression depending on its spatial niche context. In CSN-R4, the tumor-immune interaction niche, Keratin+ tumor cells exhibited upregulation of HLA-Class 1 and HLA-DR compared to those in other niches, indicating enhanced antigen-presenting capacity^55,56^ (Fig. 5f, Supplementary Fig. 16c). This may facilitate tumor antigen recognition by CD8+ T cells, which were also enriched within this niche, thereby promoting immune surveillance and cytotoxic clearance. These tumor cells also expressed elevated levels of Vimentin and α-SMA, canonical markers of EMT, suggesting the acquisition of partial mesenchymal features^57,58^ (Fig. 5f, Supplementary Fig. 16c). Enriched macrophage populations were observed in CSN-R2 and CSN-R4 (Fig. 5a), where they exhibited higher expression of HLA-DR, CD11c, and CD45 (Fig. 5g, Supplementary Fig. 16d), consistent with a highly antigen-presenting and activated phenotype^57^. This state reflects their proinflammatory potential and functional role as key effectors within an anti-tumor immune microenvironment^59^. NNC detection revealed that the CSN-R4 was in spatial proximity to two distinct niches (Fig. 5h). One was BCN-4, the major tumor niche composed of large aggregates of Keratin+ tumor cells. The other was CSN-R2, the macrophage dominated immune niche. This spatial co-localization pattern was observed in both the mixed and compartmentalized subtypes, with greater prevalence in the compartmentalized group (Supplementary Figs. 14 and 15). Further analysis showed that a higher proportion of CSN-R2 was significantly associated with improved patient prognosis (Fig. 5i, Supplementary Figs. 17a). In contrast, stratifying samples based solely on the total proportion of macrophages across the entire tissue failed to demonstrate a similar prognostic association (Supplementary Figs. 17b). These observations suggest that the presence of structured macrophage-rich niche, rather than merely the abundance of macrophages, plays a critical role in shaping effective anti-tumor immune responses and underscores the advantage of niche-based analysis over cell-level approaches.

### Harmonics delineates structured immune niches and resolves spatial functional divergence in colorectal cancer

We applied Harmonics to a CRC dataset profiled by CODEX (CODEX CRC)^5^, which comprises two subtypes. The Crohn’s-like reaction (CLR) group is characterized by the formation of numerous tertiary lymphoid structures^60^ (TLSs) at the tumor invasive front and is associated with improved patient survival, whereas the other group lacks TLSs and instead displays diffuse inflammatory infiltration (DII). We designated the DII group as the control and applied Harmonics to identify CSNs in the CLR group (Supplementary Fig. 18a to d). As expected, Harmonics consistently detected TLSs by revealing two spatially co-localized CSNs corresponding to the B cell zones (CSN-R0) and T cell zones (CSN-R1), capturing distinct spatial compositions within a unified immune structure (Fig. 6a, Supplementary Fig. 18e). In contrast, although some baseline methods partially included TLSs within CSNs, none were able to resolve the substructures of the B cell and T cell zones.

**Fig. 6.**
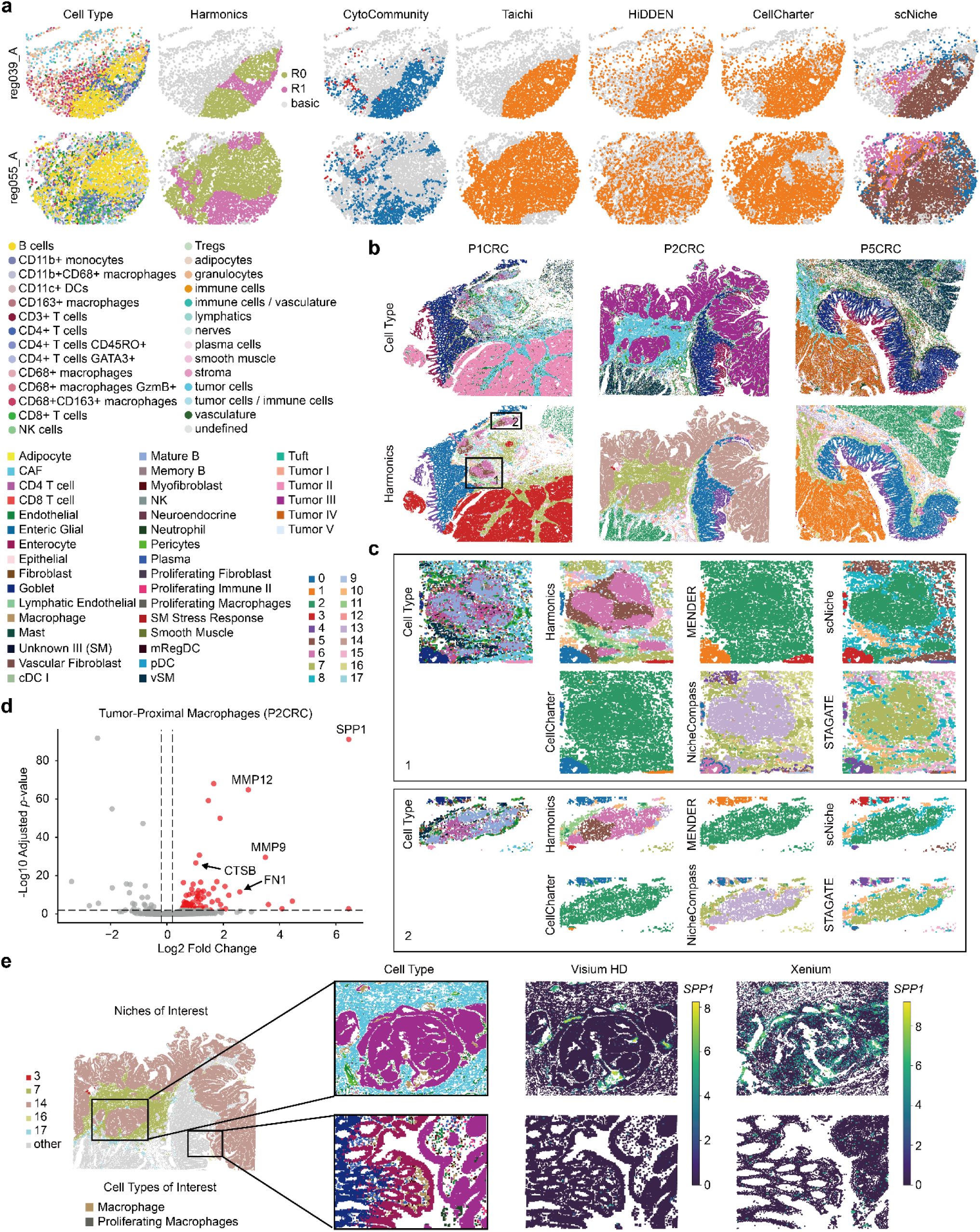
Harmonics deciphers immune architecture and spatially variable gene expression in colorectal cancer. **a,** two representative slices from the CLR group of the CODEX CRC dataset, shown with cell type annotations and CSNs identified by Harmonics and baseline methods. CSNs identified by Harmonics are labeled with the prefix "R", and "basic" denotes niches not recognized as CSNs. **b,** the three slices of the Visium HD CRC dataset, colored by cell type and CN assignments defined by Harmonics. Two representative ROIs focusing on TLSs are outlined. **c,** comparison of CN assignments from Harmonics and baseline methods within the two ROIs. Only Harmonics successfully distinguishes B cell zones from T cell zones. **d,** volcano plot of differentially expressed genes in tumor-proximal macrophages from the P2CRC slice. Genes with an adjusted *p*-value below 0.01 and log2 fold change greater than 0.2 are considered significantly upregulated and are highlighted in red. **e,** expression patterns of the representative upregulated gene *SPP1* in tumor-proximal macrophages from the P2CRC slice. Both Visium HD and the corresponding Xenium slices are shown. The full tissue slice is colored by tumor and nearby niche annotations, with all other niches shown in light gray. Zoomed-in ROIs are colored by cell types and normalized *SPP1* expression levels.

We further applied Harmonics to a Visium HD CRC dataset consisting of three CRC slices (Visium HD CRC)^9^, namely P1CRC, P2CRC, and P5CRC. This dataset was analyzed at near single-cell resolution using 8-μm bins filtered for predicted singlets (Methods). Harmonics automatically determined 18 CNs and revealed strong consistency across slices. For instance, cancer associated fibroblasts (CAF)-rich regions adjacent to tumors were consistently identified as CN-7, goblet cell-enriched niches as CN-0, and enterocyte-dominated regions as CN-4. Moreover, Harmonics distinguished tumor subtype-specific CNs (CN-1, CN-3, and CN-14), each predominantly present in one slice (Fig. 6b, Supplementary Fig. 19a to c). Notably, Harmonics preserved its ability to delineate the B cell and T cell zones of TLSs, while the condition-agnostic baseline methods also either missed TLSs entirely or were unable to resolve their substructures (Fig. 6c, Supplementary Fig. 20). These findings highlight the ability of Harmonics to accurately identify structured immune architectures and capture spatial heterogeneity within the tumor microenvironment.

Subsequently, we focused on macrophages within the Visium HD CRC dataset and categorized them into tumor-proximal and tumor-distal populations (Supplementary Fig. 21a). Specifically, based on NNC patterns (Supplementary Fig. 19d), we found that all three tumor niches exhibited spatial proximity to two non-tumor CNs, CN-7 and CN-16, which represented CAF-dominant and neutrophil-enriched microenvironments, respectively. In addition, CN-17 was primarily localized adjacent to CN-7 and was simultaneously enriched for both CAFs and macrophages, suggesting a mixed niche associated with CAF-rich regions. Based on these spatial relationships, macrophages located in CN-7, CN-16, CN-17, and the three tumor CNs were classified as tumor-proximal, while those in all other niches were considered tumor-distal. Differential expression analysis revealed spatially distinct gene expression patterns in macrophages, with tumor-proximal macrophages in CRC exhibiting a profile characteristic of an immunosuppressive, M2-like phenotype^61^ (Methods). These macrophages typically showed upregulation of *SPP1*, which has been shown to promote colorectal cancer cell migration and invasion by enhancing EMT and modulating tumor-stroma interactions^62,63^ (Fig. 6d, Supplementary Fig. 21b). They also expressed elevated levels of *MMP9* and *MMP12*, matrix metalloproteinases that degrade extracellular matrix components and are often co-expressed in spatial proximity to CAFs, suggesting coordinated interactions that facilitate extracellular matrix remodeling and local tumor progression^64,65^. This expression pattern reflects a protumoral macrophage state that supports tissue remodeling, invasion, and immune evasion within the tumor-proximal microenvironment. Further validation using both Visium HD and paired Xenium datasets confirmed that the functional state of macrophages is closely associated with their spatial context, and demonstrated that Harmonics is capable of capturing such spatially contextualized CN structures (Fig. 6e, Supplementary Fig. 21c).

## Discussion

In this paper, we presented Harmonics, a computational framework for comprehensive characterization of both common and condition-specific cell niches in spatial omics data. Harmonics centers on the HDM algorithm, which relies on the similarity assumption between the cell type distributions of each niche and that of the local neighborhoods of its constituent cells. This strategy enables robust, efficient, and scalable niche identification, further enhanced by downstream modules implemented in Harmonics for detailed niche characterization. Harmonics operates in two modes. For condition-agnostic datasets, it identifies CNs directly across the entire dataset. We demonstrated its superior accuracy and ability to identify fine-grained structures compared to baseline methods across datasets spanning diverse technologies and scales. In case-control settings, Harmonics first defines BCNs in the control group and then delineates CSNs in the case group using the BCNs as a reference. We showcased the ability of this framework in disease-related applications by uncovering disease-associated dynamics, subtype-specific spatial patterns, and structured immune niches in three representative diseases including PF, TNBC, and CRC. Unlike most existing approaches, Harmonics uses cell type annotations as input rather than directly relying on raw molecular features, enabling seamless integration with diverse spatial omics platforms and ensuring strong generalizability. In contrast to the existing methods, Harmonics is capable of explicitly identifying multiple CSNs by incorporating condition-specific information of samples. Collectively, these features make Harmonics a highly practical and generalizable tool for spatial niche analysis.

Harmonics has its limitations that warrant further improvement. First, since Harmonics relies on cell type annotations as input, the quality and granularity of the resulting niche assignments are inherently constrained by the resolution and accuracy of these annotations. This dependence requires users to have a certain level of understanding of their specific biological question and to apply appropriate tools for cell clustering or annotation. Our experiments have demonstrated that cell type annotations provided in most studies are of sufficient quality to support effective niche identification. Second, the current definition of cell neighborhood in Harmonics treats all neighboring cells as equally important when calculating the local cell type distribution. This uniform weighting does not account for the spatial distances between a cell and its neighbors, which may lead to less precise delineation of niche boundaries. A potential improvement would be to incorporate multi-range neighborhood definitions that reflect the relative proximity of surrounding cells. Third, the current method for automatically determining the number of niches in Harmonics, although effective, still relies on a pre-defined similarity threshold. While this threshold was derived from annotated datasets and has shown good performance in our experiments, developing a fully data-driven approach that avoids the need for threshold selection would be a valuable future direction. However, owing to the hierarchical nature of the HDM algorithm, Harmonics retains stable niche configurations across a range of niche numbers, thereby enabling users to directly compare results at varying levels of granularity without re-running the algorithm.

Several functional extensions may further enhance the utility of Harmonics in future applications. First, emerging methods that aim to directly infer cell type labels from high-resolution H&E-stained images^66,67^ may, once matured and validated, allow Harmonics to define niches based on image-derived cell type information. This would substantially reduce experimental costs and expand applicability. Second, incorporating gene regulatory information, such as ligand-receptor interaction networks into the downstream module for niche characterization may enrich biological interpretation by providing insights at the pathway or regulatory level, thus enabling finer-grained resolution of underlying processes^19^. Third, when applied to serial tissue sections with spatial continuity, pre-alignment of slices could be performed to enable construction of three-dimensional (3D) cell neighborhood structures^30^. This would enable the identification of 3D niches and support more comprehensive characterization of spatial tissue architecture.

In summary, Harmonics holds strong promise as a practical and versatile tool for spatial niche analysis due to its robustness, efficiency, scalability, interpretability, and generalizability, along with a rich suite of downstream modules. We envision it to be a foundational method for niche analysis across diverse biological and technological contexts.

## Methods

Harmonics is an accurate and robust computational framework designed to characterize niches in spatial omics datasets. It is originally developed for datasets with single-cell or near single-cell resolution, and requires only the spatial coordinates of each cell and a corresponding predefined cell type or cluster identity. To broaden its applicability, we further extend Harmonics to work on low-resolution (i.e., spot-level) spatial omics data, as long as the cell type deconvolution results are provided. In addition to the core model of Harmonics, we also implement a set of downstream modules, including cell type enrichment analysis, potential CCI detection analysis, and NNC pattern analysis, to support further biological interpretation.

### The model of Harmonics

Harmonics supports two operational modes. First, for the condition-agnostic datasets, Harmonics identifies CNs using HDM directly from the slices and determines the optimal number of CNs depending on the minimum difference between any pairs of CNs. Second, under case-control settings, Harmonics assumes that the case group not only shares some common CNs with the control group, but also comprises some CNs that are specific to the case group. In this setting, Harmonics first defines a set of BCNs from the user-defined control group, and then identifies CSNs in the case group using the BCNs as a reference, while also determining the number of CSNs automatically.

Since the procedure for defining BCNs in the control group is identical to the procedure used in the condition-agnostic setting, we are going to elaborate our method directly under the case-control setting.

We denote the data form the control group as *D*^ctr^, consisting of *S*^ctr^ slices and a total of 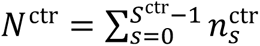 cells, where 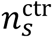 represents the number of cells in the *s*-th slice of the control group. For each slice *s*, let 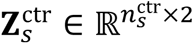 denote the matrix of spatial coordinates, where each row corresponds to one cell. We define 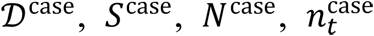 and 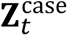 analogously for the case group, where *t* is the slice index for the case group.

### Cell neighborhood construction

For each cell, we represent it using the cell type distribution of its neighborhood. Specifically, for each slice, we treat all cells as vertices and construct edges between them using Delaunay triangulation^68^ based on 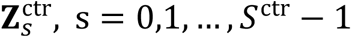 and 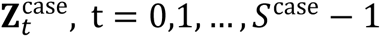. To avoid the influence of spurious long-range connections, we remove the top 1% longest edges. This results in an adjacency graph for each slice. Based on these adjacency graphs, we define the cell neighborhood of each cell as the set of cells (including itself) that are reachable within *h* hops. However, after the removal of long-range edges, some cells may have only a limited number of neighboring cells within *h* hops. To ensure that each cell has a well-defined neighborhood, we apply a correction: for any cell whose neighborhood contains fewer than *l* cells, we augment the neighborhood by including the closest cells that are not already in it, until the total number of cells reaches *l*. We then count the number of cells of each cell type within this neighborhood, obtaining a *K* - dimensional vector for each cell, where *K* is the total number of cell types in the dataset. This *K* -dimensional vector is then normalized to obtain the cell type distribution within the neighborhood, which serves as the final representation of the cell. As a result, we obtain two representation matrices, denoted as 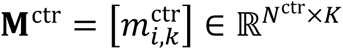 for the control group and 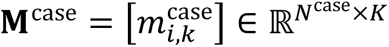 for the case group, where each row corresponds to the cell type distribution of the neighborhood of one cell. By default, we set *h* = 3 and *l* = 20 unless stated otherwise.

### Delineation of BCNs

Harmonics adopts HDM as its core strategy to obtain a series of stable solutions with varying numbers of niches. We begin by applying k-means clustering to **M**^ctr^, using a user-defined number of clusters *Q*_max_, which is intentionally set to be apparently larger than the estimated optimal number of BCNs in the control group. For each cluster (i.e. initial niche), Harmonics computes the number of cells of each cell type and normalizes this count to obtain a *K*-dimensional cell type distribution vector. Collectively, these vectors form a matrix 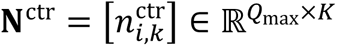, where each row corresponds to the cell type distribution of one predefined BCN.

Harmonics assumes that if a cell truly belongs to a specific niche, then the cell type distribution of its local neighborhood should closely resemble the overall cell type distribution of that niche. Conversely, if the neighborhood distribution of the cell deviates substantially from that of a given niche, the cell is likely to be involved in distinct biological processes and therefore should not be assigned to that niche.

Based on this assumption, for each cell in the control group, Harmonics evaluates the divergence between its neighborhood and each of the BCNs by comparing their cell type distributions. This yields a divergence matrix 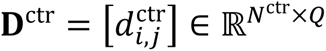, where 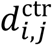 denotes the dissimilarity between the *i*-th cell neighborhood in the control group and the *j*-th BCN, and *Q* represents the current number of BCNs. We use JSD^25,26^ as the metric to quantify this dissimilarity which is calculated as:

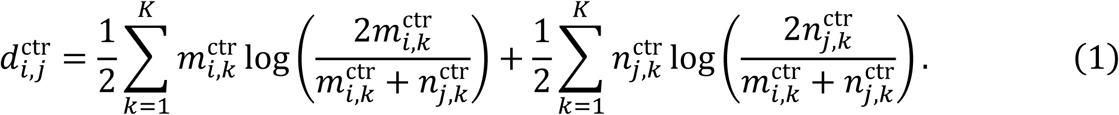

Each cell is assigned to the BCN whose cell type distribution is closest to its neighborhood, i.e., the niche corresponding to the minimum entry in the respective row of **D**^ctr^. Next, the cell type distribution of each BCN is re-calculated based on the current assignment of cells to niches, leading to an updated **N**^ctr^. Harmonics then iteratively alternates between assigning cells to BCNs and updating the corresponding niche distributions. This process continues until the change in **N**^ctr^ between two consecutive iterations, measured by the Frobenius norm, falls below a convergence threshold *tol*. To address situations where the number of niches is inappropriate for the data and the algorithm fails to converge, we set the maximum number of iterations to 100. Through this procedure, we ultimately obtain an assignment of each cell to one of the BCNs under the current number of niches and the cell type distribution of each BCN.

There are two main reasons for choosing JSD. First, compared to Kullback-Leibler Divergence (KLD), JSD is a symmetrized and smoothed version with a bounded range, which results in to more stable and interpretable comparisons. Second, unlike standard geometric distances such as Euclidean distance, JSD is better suited for capturing differences between probability distributions. We validated that using JSD achieves better trade-offs between accuracy and robustness compared to other commonly used metrics (Supplementary Fig. 2).

Subsequently, Harmonics measures the divergence between each pair of BCNs using WJSD, resulting in a symmetric matrix 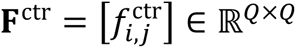, where 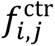 denotes the WJSD score between the cell type distributions of the *i*-th and *j*-th BCNs. Here 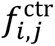 is set to +∞ if *i* = *j*; otherwise, it is computed as follows:

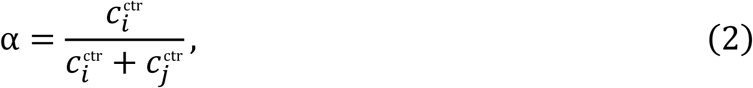

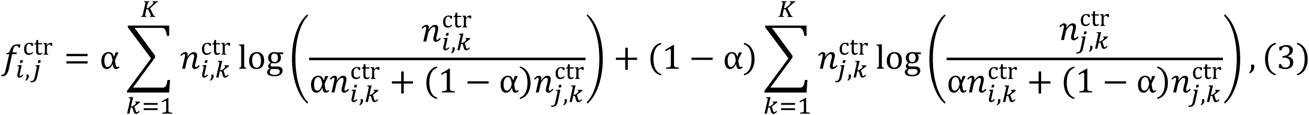

where 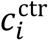 and 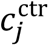 denote the number of cells assigned to the *i* -th and *j* -th BCNs, respectively. Equations (2) and (3) can be interpreted as the average KLD between the neighborhood cell type distributions of all cells in the *i* -th and *j*-th BCNs and the true distribution of the merged niche, while approximating each neighborhood distribution by the cell type distribution of the niche to which the cell is assigned. This design allows Harmonics to balance between the similarity of the two niche and the relative sizes of the niches involved. Afterward, Harmonics merges the pair of BCNs with the smallest divergence score, as indicated by the minimum entry in **F**^ctr^, thereby reducing the number of BCNs by one. Harmonics then iteratively performs niche merging and re-optimizes cell-to-niche assignments under the current number of niches, until the number of BCNs reduces to a user-defined lower bound *Q*_min_. The above procedure is referred to as HDM.

After obtaining a series of solutions with varying numbers of niches, Harmonics calculates the pairwise JSD between all niches within each solution. For each candidate niche number, it uses the minimum JSD among all niche pairs to quantify the similarity between the most similar niche pairs, and denotes this value as the *minJSD* score for that solution. Although the *minJSD* score is not strictly monotonic, it generally increases as the number of niches decreases. Harmonics uses the *minJSD* score as an indicator and defines the optimal solution as the point where the curve first crosses a predefined threshold *β*^ctr^. This criterion captures the moment when the most similar niches become sufficiently distinct. Unlike metrics that depend on external clustering techniques, this strategy is inherently integrated in the Harmonics framework, ensuring strong methodological consistency and interpretability. With an appropriately chosen *β*^ctr^, this approach avoids over-clustering, where highly similar niches are unnecessarily split, as well as under-clustering, where biologically distinct niches are incorrectly merged.

In summary, Harmonics ultimately produces a cell type distribution matrix **N**^ctr*^ = 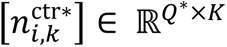 along with the niche assignment label for each cell in the control group, where *Q*^∗^ denotes the optimal number of BCNs identified. In settings where niche detection is performed on datasets comprising a single group only, this yields the final niche assignment.

In practice, we calibrate the threshold *β*^ctr^ using two annotated datasets, STARmap V1C^27^ and MERFISH FC&S^1^, by selecting the value of *β*^ctr^ such that the number of niches automatically determined by Harmonics basically matches the number of annotated niche categories in these datasets. Based on this calibration, we set *β*^ctr^ = 0.1 and kept it for all experiments throughout this study. Additionally, we fix *tol* = 1 × 10^−4^ and *Q*_min_ = 2 across all experiments. The maximum number of initial niches, *Q*_max_, is theoretically dataset-dependent; by default, we set *Q*_max_ = 20, and further details on its selection will be discussed later.

### Identification of CSNs

Harmonics assumes that data from the case group contain both BCNs common to the control group and a number of CSNs unique to the case group. Accordingly, Harmonics uses the previously determined **N**^ctr∗^ as a reference to guide the discovery of CSNs.

To initialize the process, for each cell in the case group, Harmonics computes the JSD between its neighborhood cell type distribution and that of each BCN. If the minimum JSD is below a predefined threshold *β*^init^, it confidently assigns the cell to the corresponding BCN. The remaining unassigned cells are initialized into *R*_max_ clusters using k-means clustering based on their representations in **M**^case^, where *R*_max_ is deliberately set to exceed the estimated number of CSNs. Harmonics then recomputes the cell type distributions for these newly initialized CSNs to form the matrix 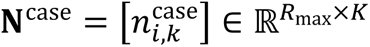, while keeping the distributions of BCNs fixed.

Subsequently, Harmonics applies HDM to the case group to obtain a series of niche assignment results under varying numbers of CSNs. For a given CSN count *R*, each cell is assigned to the niche (either a BCN or CSN) with the smallest JSD relative to its neighborhood distribution. During this process, only the CSN distributions in **N**^case^ are updated, while the BCNs remain fixed. The convergence criterion is set the same as that used in BCN discovery.

In the merging phase, each iteration allows for either merging two CSNs or merging of a CSN into a fixed BCN. The similarity between niches is quantified using WJSD. To avoid bias arising from the potentially low abundance or absence of certain BCNs in the case group, Harmonics assumes equal cell counts when comparing a CSN with a BCN, which causes WJSD to degrade to the standard JSD while preserving its numerical range^26^. Harmonics compares all pairwise WJSD values and merges the most similar niche pair. Notably, even when a CSN is merged into a BCN, the original cell type distribution of the BCN remains unchanged.

This hierarchical merging process continues until all CSNs have been absorbed into the fixed BCNs, resulting in zero remaining CSNs. To determine the optimal number of CSNs, we adopt the same strategy used in selecting the optimal number of BCNs, identifying the point where the *minJSD* score first crosses the threshold *β*^case^.

Finally, in the case group, Harmonics outputs the cell type distribution matrix for all CSNs, denoted as 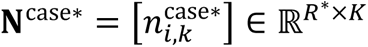, along with a niche assignment label for each cell, indicating the specific niche (either one of the *R*^∗^ identified CSNs or to a fixed BCN) to which the cell is assigned.

We fix *R*_max_ = 10 across all experiments, and set *β*^init^ = *β*^case^ = 0.1, which is consistent to *β*^ctr^.

### Niche discovery in low-resolution data

Harmonics can be extended to identify CNs and CSNs in low-resolution spatial omics datasets. In technologies such as Visium^69^, each spot on the sequencing array typically comprises multiple cells, making single-cell resolution unattainable. To address this, various methods have been developed for cell type deconvolution^30,70,71^, which estimate the proportions of different cell types within each spot. Harmonics can leverage these deconvolution results in place of single-cell labels to enable niche identification. For low-resolution data, Harmonics represents each spot and each niche by averaging the cell type proportions of neighboring spots and assigned spots, respectively. All other steps remain consistent with those applied to datasets with single-cell labels.

### Facilitating niche definition via label transfer

For large-scale datasets with minimal inter-slice heterogeneity, such as serial sections from the same tissue, Harmonics supports a label transfer strategy to improve computational efficiency. Specifically, the dataset can be split into two groups at the slice level. Harmonics defines CNs using one group and assign cells in the other group to the most similar CN based on their neighborhood cell type distributions. In this study, unless otherwise stated, we apply Harmonics directly to the full dataset without employing label transfer.

### Niche representation refinement

When the annotated cell types include many that are of very low abundance, have limited functional significance, and are irrelevant to the spatial patterns of interest, Harmonics allows for refinement of niche representations to stabilize the overall procedure. This refinement emphasizes the contribution of important cell types while reducing the influence of less informative ones. Specifically, when updating the cell type distribution for each niche, Harmonics considers only the top *K*^re^ most abundant cell types in that niche. We do not apply this refinement in this study unless otherwise specified.

### Downstream modules for niche characterization

#### Cell type enrichment test

For a given dataset, Harmonics supports cell type enrichment analysis based on the assigned niche labels, aiding the biological interpretation of identified niches. Specifically, for each niche, Harmonics calculates the number of cells from each annotated cell type and conducted a two-sided Fisher’s exact test^72^ to evaluate whether the proportion of a given cell type in the niche significantly deviates from its proportion in the remaining niches, which serve as the background. The *p*-values obtained are adjusted using the Benjamini–Yekutieli (B-Y)^73^ procedure to control the false discovery rate (FDR). The log2 fold change, denoted as *Log*2*FC*_ct_, is computed as the log2 ratio between the proportion of a given cell type within the tested niche and its proportion across the background niches. In our study, a cell type is considered enriched in a given niche if it exhibits an adjusted *p*-value less than 0.05, a *Log*2*FC*_ct_ greater than 1, and accounts for more than 1% of the cells in that niche.

#### Potential CCI detection

Harmonics detects potential CCIs by assessing the enrichment of spatially adjacent cell type pairs within each niche. Specifically, for a given dataset, Harmonics constructs cell adjacency graphs for each slice using Delaunay triangulation and remove the top 1% longest edges to define spatial proximity. To test whether the adjacency of a specific cell type pair is enriched in a given niche, Harmonics treats all other niches as background and performs a two-sided Fisher’s exact test. The test evaluates whether the proportion of edges connecting the cell type pair within the niche significantly deviates from their proportion in the background. FDR is controlled using the B–Y procedure. The log2 fold change, denoted as *Log*2*FC*_cc_, is defined as the log2 ratio between the proportion of a specific type of edge within the tested niche and its proportion in the background. In this study, a cell type pair is considered to exhibit a potential CCI within a niche if the corresponding edges account for more than 1% of all edges in that niche, the adjusted *p*-value is below 0.05, and *Log*2*FC*_cc_ exceeds 1

#### NNC pattern identification

Harmonics identifies NNC patterns by evaluating whether a specific target niche is more likely to localize in the neighborhood of a given source niche compared to other niches. Harmonics first constructs cell–cell adjacency relationships in the similar manner as in CCI detection, but treats all edges as bidirectional. A two-sided Fisher’s exact test is then performed to assess whether there is a significant deviation between the following two proportions: (i) the proportion of edges that originate from a specific source niche and terminate at a given target niche, relative to all edges originating from that source niche; and (ii) the proportion of edges that originate from all other (non-source) niches and terminate at the same target niche, relative to all edges originating from non-source niches. Edges connecting cells within the same niche are excluded from the analysis. The log2 ratio between the two proportions is denoted as *Log*2*FC*_nn_, and the B-Y procedure is applied to control the FDR. A target niche is considered co-localized with a source niche if the adjusted *p*-value is below 0.05, *Log*2*FC*_nn_ exceeds 1, and the value of proportion (i) is greater than 0.01.

### Data description

#### STARmap V1C dataset

The STARmap V1C dataset^27^ consists of a single slice of mouse V1 cortex, containing 1,207 cells and 1,020 genes. It includes annotations of seven distinct niches and 16 cell types. For benchmarking purposes, we excluded the smooth muscle cell ("Smc") population, which represents a small number of epidermal cells that are coarsely annotated as layer 1 due to their limited abundance.

#### MERFISH FC&S dataset

The MERFISH FC&S dataset^1^ includes slices of mouse frontal cortex and striatum from three developmental periods: 4-week-old (10 slices), 24-week-old (8 slices), and 90-week-old (13 slices), with spatially resolved expression profiles of 374 genes. We excluded regions labeled as "brain ventricles" due to their extremely low representation (less than 1% of total cells), resulting in a final cell count of 123,278, 89,985, and 162,844 cells for the three groups, respectively. Cells were categorized into 43 distinct cell types and annotated into seven major niches.

#### Visium DLPFC dataset

The Visium DLPFC dataset^29^ is a low-resolution, spot-level SRT dataset comprising slices of the human dorsolateral prefrontal cortex from three human donors. Each donor group contains four tissue slices. For each group, we only retained genes detected in all slices within the group and removed mitochondrial-encoded genes. After filtering, group 1 (samples 151507 to 151510) consists of 17,985 spots and 18,938 genes, group 2 (samples 151669 to 151672) contains 15,101 spots and 19,177 genes, and group 3 (samples 151673 to 151676) includes 14,243 spots and 19,551 genes. Expert annotations defined seven niches for groups 1 and 3, and five niches for group 2.

### MERFISH WMB dataset

The MERFISH WMB dataset^31^ consists of 23 consecutive sagittal slices of a mouse hemisphere, comprising a total of 2,081,549 cells and 1,122 genes, categorized into 338 cell types. The dataset is coarsely divided into 15 major brain regions, including one undefined region.

### Xenium PF dataset

The Xenium PF dataset^2^ consists of two groups of human lung tissue slices: one from healthy donors and the other from patients with pulmonary fibrosis. Each slice includes a visually estimated pathology percentage annotated by experts, indicating the extent of remodeling within the tissue. We excluded one healthy sample (THD0011) whose pathology percentage exceeded that of some disease samples. After filtering, the dataset includes 9 healthy slices (199,773 cells) and 35 disease slices (1,405,182 cells), with 343 genes profiled across all samples. Cells are categorized into 47 cell types.

#### MIBI-TOF TNBC dataset

The MIBI-TOF TNBC dataset^12^ consists of one tumor slice per patient from 40 individuals with TNBC. Based on the abundance and spatial organization of immune cells, the slices are categorized into three groups: cold group (6 slices), with few immune cells; compartmentalized group (15 slices), where immune cells and tumor cells are spatially separated; and mix group (19 slices), where immune and tumor cells are intermixed. Survival data is available for all individuals except for patient 22. For analysis, we designated the cold group as the control group, and combined the compartmentalized and mix groups as the case group. An undefined cell type ("Unidentified") was excluded from the analysis, resulting in 22,748 cells in the control group and 173,205 cells in the case group, with 36 proteins profiled across all samples. A total of 17 cell types were annotated.

#### CODEX CRC dataset

The CODEX CRC dataset^5^ comprises CRC tissue slices from two subtypes, namely the DII group and the CLR group. We designated the DII group as the control group and the CLR group as the case group. Three slices (reg023_A, reg064_A, and reg066_A) with more than 50% of immune cells not clearly identified were excluded from the analysis, resulting in 72 control slices (143,062 cells) and 65 case slices (103,274 cells), with 56 proteins profiled across all samples. Cells were annotated into 28 distinct cell types.

#### Visium HD CRC dataset

The Visium HD CRC dataset^9^ includes three CRC tissue slices, with cell type deconvolution performed on 8-μm bins. Only bins predicted as singlets were retained for analysis, resulting in a total of 810,099 bins, 18,085 genes, and 38 cell types. Each slice is also accompanied by paired SRT data profiled using the Xenium platform.

### Benchmarking analysis

For the task of identifying CNs in condition-agnostic datasets, we compared Harmonics with several baseline methods, including MENDER^28^, scNiche^18^, CellCharter^17^, NicheCompass^19^, the unsupervised mode of CytoCommunity^24^, STAGATE^15^, and GraphST^16^.

For the detection of the CSNs in the case-control settings, baseline methods specifically designed for this task include the supervised mode of CytoCommunity, Taichi^22^, and HiDDEN^23^. In addition, we included CellCharter and scNiche for comparison, as both methods provide modules to automatically determine the optimal number of niches. However, CytoCommunity, CellCharter, and scNiche do not directly associate the identified niches with condition labels. To enable a broader comparison, we adopted a relaxed definition of CSNs for these methods: any niche whose proportion in the case group is higher than in the control group is considered disease-relevant or condition-relevant and therefore included for comparison.

For datasets with niche annotations, all methods were configured to output the same number of niches as the annotated classes. For other datasets, methods with automatic cluster number selection used their own estimated optimal solution, while for methods without this functionality, the number of niches was set to match that determined by Harmonics.

#### Harmonics

For the Xenium PF dataset, due to its highly non-convex tissue architecture, we set *h* = 2 when constructing cell neighborhoods. For the Visium DLPFC dataset, since each spot represents a mixture of multiple cells rather than single-cell resolution, we also set *h* = 2. We performed cell type deconvolution using our previously developed method, STitch3D^30^, using the same procedure described in the online tutorial, and represented each spot by the proportions of 18 cell types. For group 1 and group 2 of the Visium DLPFC data, we set *K*^re^ = 10 and *K*^re^ = 5, respectively, for niche representation refinement. For the MERFISH WMB dataset, which spans the entire mouse brain hemisphere and is expected to contain a large number of niches, we initialized the maximum number of niches *Q*_max_ to 100. For the Visium HD CRC dataset, the default setting of *Q*_max_ = 20 led to a warning indicating that no suitable solution was found. Therefore, we increased *Q*_max_ to 30. All other parameters were kept consistent with the default settings described in the section "The model of Harmonics".

#### MENDER

MENDER identifies CNs based on cell type annotations. It constructs multi-scale neighborhoods around each cell, with the *radius* parameter specifying the distance between two adjacent scales. As this parameter is dataset-dependent, we set it to 150 for the STARmap V1C dataset, 0.5 for the Visium HD CRC datasets, and 15 for other datasets. The target number of niches was set to match the number identified by Harmonics, and MENDER’s built-in resolution search was used to obtain the corresponding solution. All other parameters were kept at their default values, and the implementation followed the online tutorial.

#### scNiche

scNiche adopts a multi-view clustering framework^74^ to integrate cell type information and profiled biological features, and can automatically determine the optimal number of niches. For all SRT datasets except STARmap V1C, we first applied PCA to reduce the dimensionality of the gene expression matrix to 50, which was then used as input for scNiche. To reduce memory usage, scNiche offers a batch training strategy. Following the online tutorial, we set the number of batches to 2, 20, and 100 for the Visium DLPFC dataset, the MIBI-TOF TNBC dataset, and the MERFISH FC&S dataset, respectively. For the large-scale datasets MERFISH WMB and Xenium PF, both containing over 1.5 million cells, we set the number of batches to 200. Since STARmap V1C dataset includes only a single slice, batch training was not used. For the remaining datasets, the number of batches was set to 100. In the case of the Visium DLPFC dataset, where scNiche requires cell type deconvolution results as input, we used the output generated by STitch3D, consistent with that used in Harmonics. For the automatic selection of optimal solution, we used the same search range for the number of niches as in Harmonics to ensure a fair comparison. All other parameters were kept at their default values, and the implementation followed the online tutorial.

#### CellCharter

CellCharter builds spatially informed representations based on embeddings of cells obtained from existing single-cell integration methods, and automatically determines the optimal number of niches. Following its official tutorial, we used scVI^75^ and trVAE^76^ to obtain embeddings for SRT data and spatial proteomics data, respectively. The search range for the number of niches during automatic selection was set to match that used by Harmonics. All other parameters were kept at their default values, and the implementation followed the online tutorial.

#### NicheCompass

NicheCompass characterizes spatial cell niches by learning communication-aware cell embeddings that capture local signaling programs across spatial omics datasets, and includes a covariate embedding module to tackle batch effects. We set the dimensionality of the covariate embedding to the minimum of the number of slices and 10. Since the NicheCompass procedure uses Leiden clustering to determine niche assignments but does not include an automatic module for selecting the number of niches, and Leiden clustering does not allow direct specification of the number of clusters, we applied the Gaussian mixture model (GMM) from the scikit-learn package^77^ to cluster the learned embeddings into the same number of niches as determined by Harmonics. All other parameters were kept at their default values, and the implementation followed the online tutorial.

#### CytoCommunity

CytoCommunity identifies niches using only cell type information through a GNN framework. It includes two modes. The unsupervised mode directly detects CNs within each individual slice independently, while the supervised mode incorporates condition-specific information of samples to identify disease-associated niches. The final number of niches depends in part on the training process and a majority voting step, resulting in a value less than or equal to the predefined target number. We set the target number of niches to be consistent with that used by Harmonics. In the unsupervised mode, we set the *Num*_*Run* and *Num*_*Epoch* parameters to 5 and 300, respectively, for the MERFISH FC&S dataset to reduce runtime. In the supervised mode, we set the *MineBatchSize* parameter to 32 for the MIBI-TOF TNBC dataset and 8 for the CODEX CRC dataset. All other parameters were kept at their default values, and the implementation followed the online tutorial.

#### STAGATE

STAGATE is originally designed to utilize a graph attention (GAT)^78^ autoencoder framework that integrates gene expression and spatial information to adaptively learn cell embeddings for the identification of spatial domains in SRT data. When constructing the spatial adjacency graph, we selected an appropriate radius for each dataset such that each cell had approximately 10 neighbors. The learned embedding matrix was then clustered using the mclust method to identify CNs as recommended. All other parameters were kept at their default values, and the implementation followed the online tutorial.

#### GraphST

GraphST is a GNN-based contrastive learning framework that integrates gene expression and spatial information to enable spatial clustering, batch integration, and cell type deconvolution in SRT data. For the low-resolution Visium DLPFC dataset, we set the *datatype* parameter to "10X", while for other datasets, we set the *datatype* parameter to "Slide". The embedding matrix was clustered using the mclust method to identify CNs as recommended. All other parameters were kept at their default values, and the implementation followed the online tutorial.

#### Taichi

Taichi leverages the cell representations derived from MENDER, integrates condition-specific information, and utilizes a logistic regression–based strategy to detect a single CSN within the case group. For the CODEX CRC dataset, we set the *scale* parameter to 4 and the *radius* parameter to 50, following the corresponding online tutorial. For all other experiments, we used *scale* = 6 and *radius* = 15 to match the default parameters of MENDER. All other parameters were kept at their default values, and the implementation followed the online tutorial.

#### HiDDEN

HiDDEN is a statistical framework designed to refine cell-level labels within a given condition, enabling the identification of truly condition-relevant cell populations. We used PCA for dimensionality reduction and applied the built-in method to automatically detect the optimal number of principal components. Label prediction was conducted using the logistic regression mode. All other parameters were kept at their default values, and the implementation followed the online tutorial.

### Computational efficiency

Across all datasets, we recorded the runtime of each relevant method. Harmonics consistently demonstrated high time efficiency across datasets of different scales. For condition-agnostic datasets, Harmonics was markedly faster than all baseline methods on every dataset (Supplementary Tab. 1). Under the case-control setting, Harmonics remained substantially faster than the three neural network-based methods, scNiche, CellCharter, and CytoCommunity, while performing comparably to the two logistic regression-based methods, Taichi and HiDDEN. Notably, on the CODEX CRC dataset, Harmonics even outperformed Taichi in runtime (Supplementary Tab. 2). However, due to their model design, Taichi and HiDDEN can only coarsely partition a dataset into two categories to identify a single CSN, thus addressing a highly simplified task. In contrast, Harmonics tackles a more complex task and is better suited for comprehensive CSN characterization. Time costs were evaluated using Intel^®^ Xeon^®^ Gold 5220R CPU @ 2.20 GHz (dual processor) and NVIDIA RTX A6000 48 GB GPU. Methods that supported GPU acceleration were run on GPU, while those requiring memory beyond the available GPU capacity were deemed not applicable.

### Assessment of robustness

To systematically compare robustness, each method was applied to the STARmap V1C dataset and run 20 times under different random seeds. We employed the Fowlkes-Mallows Index (FMI), a commonly used metric for assessing clustering consistency^17,18^, to evaluate the pairwise agreement between the results of different runs. A higher FMI indicates greater robustness. The same strategy was also applied when comparing different metrics used to quantify the similarity between distributions, including JSD, KLD, cross-entropy (equivalent to the reverse KLD), Manhattan distance, Euclidean distance, and cosine similarity. On the MERFISH FC&S dataset, we ran each candidate metric twice with different random seeds, and the FMI score was computed between the two runs for each slice.

For the MERFISH WMB dataset, we assessed the consistency between results obtained via label transfer and those obtained using all slices directly. In the label transfer setting, we selected one slice every four slices, starting from the first (C57BL6J-3.001), and used the selected slices to define CNs. Cells from the remaining slices were then assigned to these CNs. The FMI score was computed for each slice by comparing the CN labels in the label transfer setting with those obtained from the full dataset.

The FMI scores are computed using the implemented function in the scikit-learn (v1.6.1) package^77^.

### Quantifying niche correspondence between conditions

We first computed the cell type distribution for each niche within the control group and the case group separately. The control group contains only BCNs, while the case group includes both BCNs and CSNs. Next, for each niche pair consisting of one from each group, we calculated the JSD using logarithm base 2. We then subtracted the JSD value from 1 to obtain a similarity score ranging from 0 to 1, where higher scores indicate greater similarity in cell type composition.

### Correlation analysis

Each correlation analysis was performed between two distinct variables defined for the specific comparison of interest. Pearson’s *r* was used to quantify the strength of correlation. *P*-values were computed using a two-sided permutation test, in which the values of one variable were randomly shuffled 10,000 times. Trend lines were fitted using univariate linear regression.

### Specificity of disease-associated cell types in CSNs

In the Xenium PF dataset, we compared Harmonics with baseline methods by evaluating whether the CSNs identified by each method specifically capture disease-associated cell types. We focused on three cell types strongly linked to disease pathology. These include activated fibrotic FBs, the major fibrotic cells in fibrotic foci; transitional AT2 cells, which represent an intermediate state during the conversion of AT2 to AT1; and B cells, which are a key component of lymphoid aggregates. These three cell types are markedly more abundant in the case group than in the control group, both in absolute number and proportion. Therefore, effective CSN identification should lead to the enrichment of these cell types in specific CSNs. We define the specificity score for the *i*-th CSN as follows:

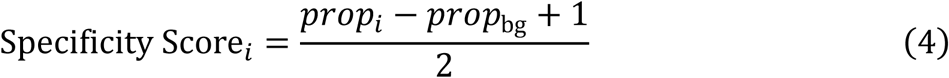

where *prop_i_* denotes the proportion of the cell type of interest within the *i*-th CSN, and *prop*_bg_ denotes its proportion in the background, defined as all other niches outside the *i*-th CSN. The score ranges from 0 to 1, with higher values indicating greater specificity. A score below 0.5 suggests that the target cell type has a lower proportion in the CSN compared to the background, which indicates poor specificity.

For each cell type, the CSN with the highest median specificity score across slices was selected for each method to enable cross-method comparison.

### Differential expression analysis

For the Xenium PF dataset, we first applied log normalization (total counts normalization followed by logarithmic transformation) to each cell’s expression profile using the Scanpy (v1.10.6) package^79^. For the MIBI-TOF TNBC dataset, since the protein expression data are already scaled per marker, no further normalization was performed.

Differential expression analysis was conducted between two groups of cells at a time. To mitigate the impact of batch effects across slices, we computed group-specific slice-level averages of the feature of interest (a gene in the Xenium PF dataset or a protein in the MIBI-TOF TNBC dataset), resulting in paired values for each slice that were used for comparison. Only slices in which both groups contained more than 20 cells were included in the analysis. *P*-values were computed using a two-sided Wilcoxon signed-rank test^80^.

For the Visium HD CRC dataset, we first removed mitochondrial genes and performed log normalization. We then classified macrophages into tumor-proximal (from CN-1, CN-3, CN-7, CN-14, CN-16, and CN-17) and tumor-distal groups (from all other CNs). Differential expression analysis was performed using the *rank_genes_groups* function in Scanpy with the Wilcoxon rank-sum test. Genes with an adjusted *p*-value below 0.01 and a log2 fold change greater than 0.2 were considered significantly upregulated.

### Projection of query data onto the reference UMAP

In the Xenium PF dataset, to compare the transcriptional states of major epithelial cells in CSN-R2 (aberrant epithelium) with those in BCN-0 (alveolar epithelium) and BCN-1 (airway epithelium), we projected CSN-R2 cells onto a reference UMAP constructed from BCN-0 and BCN-1. Four epithelial cell types were included: AT2, transitional AT2, basal, and RASC. Only slices from the case group were used with one batch-affected slice (VUILD110LA) was excluded.

The reference group contained the four cell types from BCN-0 and BCN-1, with AT2 and transitional AT2 mainly from BCN-0 (transitional AT2 being rare) and basal and RASC primarily from BCN-1. The query group contained the same four cell types from CSN-R2. Cells with fewer than 30 detected genes were removed, and the remaining cells were log-normalized. PCA was performed on the reference group using 10 components. Query cells were projected into the reference PCA space using the same loadings, after centering each gene by the corresponding mean value computed from the reference cells. Subsequently, the reference and query data were concatenated. A neighborhood graph was constructed using 15 nearest neighbors per cell and 10 principal components. UMAP embedding was then computed for visualization. All steps were performed using the Scanpy package.

### Trajectory and pseudotime inference

To investigate potential differentiation trajectories within CSN-R2 identified in the Xenium PF dataset, we applied Slingshot^81^ to infer trajectories and pseudotime for major epithelial cell types, including AT2, transitional AT2, basal, and RASC. The input embedding was based on the UMAP coordinates of cells from CSN-R2, computed as described in the section "Projection of query data onto the reference UMAP". The Slingshot model was fit for one epoch using the AT2 cluster as the root node. The analysis was performed using the pyslingshot (v0.1.5) package implemented in Python.

### Disease subtype classification by niche proportions

In the MIBI-TOF TNBC dataset, the compartmentalized and mixed groups represent two distinct disease subtypes. For both Harmonics and baseline methods, subtype labels were not accessible during niche identification. Therefore, to assess each method’s ability to capture subtype-specific spatial patterns, we evaluated how well the niche proportions could distinguish between the two subtypes. For each baseline method, we first identified the niche that best separates the compartmentalized and mixed groups. Specifically, we used the proportion of each niche across samples as input features for binary classification and computed both the AUC and its complement 1-AUC scores. All scores were pooled, and the highest score was selected to determine the most discriminative niche for each method. For Harmonics, we compared CSN-R2 and CSN-R0 against the best-performing niches from the baseline methods.

### Survival analysis

We performed survival analysis on patients from the case group of the MIBI-TOF TNBC dataset. Patient 22 lacked associated survival data and was therefore excluded, leaving a total of 33 samples for analysis. Patients were stratified into high- and low-proportion groups based on the intra-slice abundance of CSN-R2, and Kaplan-Meier survival curves^82^ were generated to compare their survival outcomes. HRs were estimated using Cox regression^83^, and the associated *p*-values were derived from a two-sided test. Stratification thresholds for CSN-R2 proportion were set to 0, 0.001, 0.005, 0.008, 0.01, and 0.03 to assess the robustness of the association across different cutoffs. To determine whether the observed survival differences were driven by niche specific effects or by dominant cell types, we repeated the analysis using the proportion of macrophages, the major cell type within CSN-R2, across the entire tissue slice. In this analysis, thresholds were selected to yield high- and low-proportion groups with patient counts matched to those in the corresponding CSN-R2-based stratifications, ensuring comparable group sizes. The analysis was conducted using lifelines (v0.30.0) package^84^.

### Evaluation metrics

We used four metrics, ARI, AMI, mF1, and wF1, to evaluate the performance of Harmonics and all baseline methods on datasets with expert-annotated niche labels. ARI and AMI quantify the similarity between two sets of label assignments based on pairwise sample agreements and mutual information, respectively, while mF1 and wF1 evaluate performance from a classification perspective by comparing predicted labels and true annotations. Before computing mF1 and wF1, we utilized the *match_cluster_labels* function provided in the online tutorial of PASTE^85^ to align the predicted niche categories from each method to the true annotations. All metrics are computed using the respective implementations provided in the scikit-learn package^77^. Their values range from 0 to 1, with higher values indicating better performance.

## Data availability

All the data used in this paper are publicly available. The STARmap V1C dataset^27^ can be accessed at [https://zenodo.org/records/7830764#.ZDpObi-1HUI]. The MERFISH FC&S dataset^1^ can be obtained from [https://cellxgene.cziscience.com/collections/31937775-0602-4e52-a799-b6acdd2bac2e]. The MERFISH WMB dataset^31^ can be downloaded from [https://cellxgene.cziscience.com/collections/0cca8620-8dee-45d0-aef5-23f032a5cf09]. The Visium DLPFC dataset^29^ is collected from [https://github.com/LieberInstitute/spatialLIBD]. The Xenium PF dataset^2^ can be obtained from GEO under accession code GSE250346. The Visium HD CRC dataset^9^ is downloaded from [https://www.10xgenomics.com/platforms/visium/product-family/dataset-human-crc]. For the spatial proteomics datasets, including the MIBI-TOF TNBC^12^ and the CODEX CRC^5^, we download it from the SODB^86^, at [https://gene.ai.tencent.com/SpatialOmics/dataset?datasetID=47] and [https://gene.ai.tencent.com/SpatialOmics/dataset?datasetID=60], respectively. All processed datasets used in this manuscript can be found at Zenodo^87^.

## Code availability

Harmonics and the tutorials for reproducing the analyses are available at https://github.com/YangLabHKUST/Harmonics. The benchmarking codes for comparison with baseline methods are deposited at Zenodo^88^.

## Acknowledgements

This work was partly supported by the Innovation and Technology Commission (ITCPD/17-9); Hong Kong Research Grants Council Grants 16301419, 16308120, 16307221, 16307322, 16302823, and 16309424; and The Hong Kong University of Science and Technology Startup Grants R9405 and Z0428 from the Big Data Institute.

## Author contributions

Y. conceived the study and supervised the project. Y. L. designed, implemented, and validated Harmonics. J. X. contributed to the experimental design. X. M. assisted with the validation of Harmonics. X. W., P. J., Z. W., and Y. C. helped with analyzing the results. Y. L. and C. Y. wrote the manuscript with input from all the authors.

## Competing interests

The authors declare no competing interests.

## Notes

### Competing Interest Statement

The authors have declared no competing interest.

