## Supplementary materials for "Hierarchical distribution matching enables comprehensive characterization of common and condition-specific cell niches in spatial omics data"

### Supplementary Tables

**Supplementary Tab. 1 | Evaluation of time cost on condition-agnostic datasets.** "NA" indicates that the method is not applicable to the corresponding dataset. For each dataset, the shortest runtime is highlighted in bold.

| Time cost (minute) | Harmonics | MENDER | scNiche | CellCharter | NicheCompass | CytoCommunity | STAGATE | GraphST |
|---|---|---|---|---|---|---|---|---|
| STARmap V1C | **0.01** | 0.97 | 0.33 | 3.08 | 13.46 | 13.51 | 0.23 | 0.44 |
| MERFISH FC&S 4-week-old group | **1.59** | 11.87 | 18.26 | 20.52 | 43.64 | 388.07 | 4.89 | 5.13 |
| MERFISH FC&S 24-week-old group | **1.44** | 13.95 | 16.38 | 20.42 | 32.75 | 258.09 | 3.84 | 4.02 |
| MERFISH FC&S 90-week-old group | **1.48** | 25.82 | 24.95 | 28.31 | 79.48 | 747.42 | 6.57 | 7.15 |
| Visium DLPFC group 1 | **0.12** | NA | 9.77 | 16.37 | 23.78 | NA | 1.45 | 1.72 |
| Visium DLPFC group 2 | **0.12** | NA | 7.44 | 14.55 | 18.13 | NA | 1.25 | 1.55 |
| Visium DLPFC group 3 | **0.21** | NA | 6.95 | 16.60 | 19.71 | NA | 1.33 | 1.53 |
| MERFISH WMB | **224.69** | 2022.35 | 1270.17 | 1475.45 | 456.79 | NA | NA | NA |
| Visium HD CRC | **8.61** | 418.52 | 351.80 | 180.00 | 601.81 | NA | 278.55 | NA |

**Supplementary Tab. 2 | Evaluation of time cost on datasets under case-control setting.** "NA" indicates that the method is not applicable to the corresponding dataset. For each dataset, the shortest runtime is highlighted in bold.

| Time cost (minute) | Harmonics | scNiche | CellCharter | CytoCommunity | Taichi | HiDDEN |
|---|---|---|---|---|---|---|
| Xenium PF | 13.83 | 667.93 | 164.68 | NA | **6.58** | 8.99 |
| MIBI-TOF TNBC | 1.51 | 96.63 | 47.72 | 1247.85 | **0.43** | 1.31 |
| CODEX CRC | 1.79 | 36.90 | 38.38 | 912.48 | 4.60 | **1.17** |

### Supplementary Figures

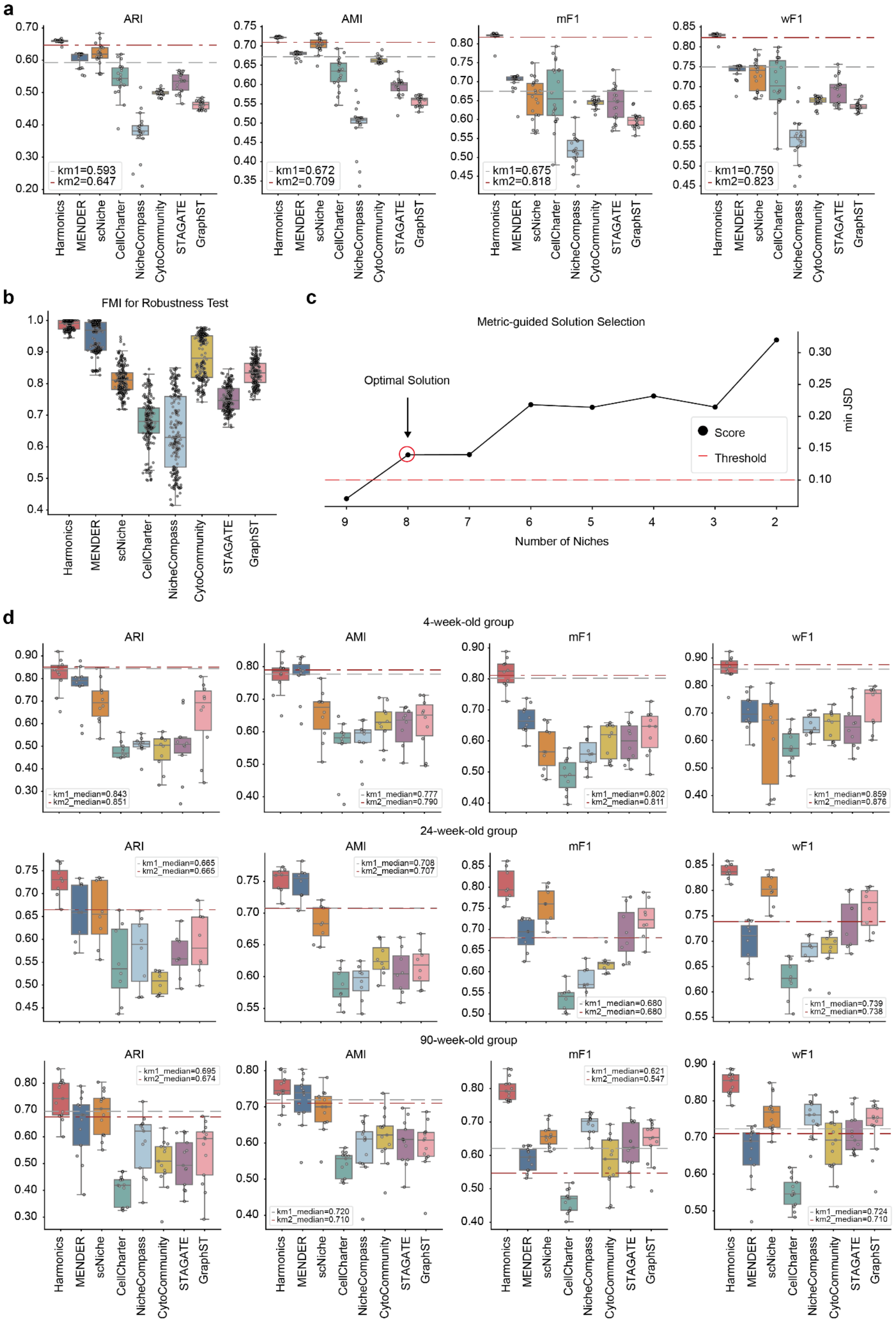

**Supplementary Fig. 1 | Performance evaluation of Harmonics and baseline methods on the STARmap V1C and the MERFISH FC&S datasets. a, d,** comparison of ARI, AMI, mF1, and wF1 scores for Harmonics and baseline methods. Each dot represents an individual run on the STARmap V1C dataset (**a**) or an individual slice from the MERFISH FC&S dataset (**d**). The gray and red dashed lines indicate two alternative approaches: the gray line shows the results of directly applying k-means to the cell representations produced by Harmonics, using the same number of clusters as annotated niches; the red line shows the results of using the HDM procedure as in Harmonics but using k-means clustering to find the solution for each niche count. Their respective scores correspond to a single run (**a**) or the median score across slices (**d**). **b,** robustness test of Harmonics and baseline methods, assessed by pairwise FMI scores across repeated runs. Each dot represents a comparison between two independent runs. **c,** $minJSD$ score curve for automatic niche number selection in the STARmap V1C dataset with smooth muscle cells included. For all box plots, the center line indicates the median; box limits represent the upper and lower quartiles; whiskers extend to the largest and smallest values within 1.5 times the interquartile range.

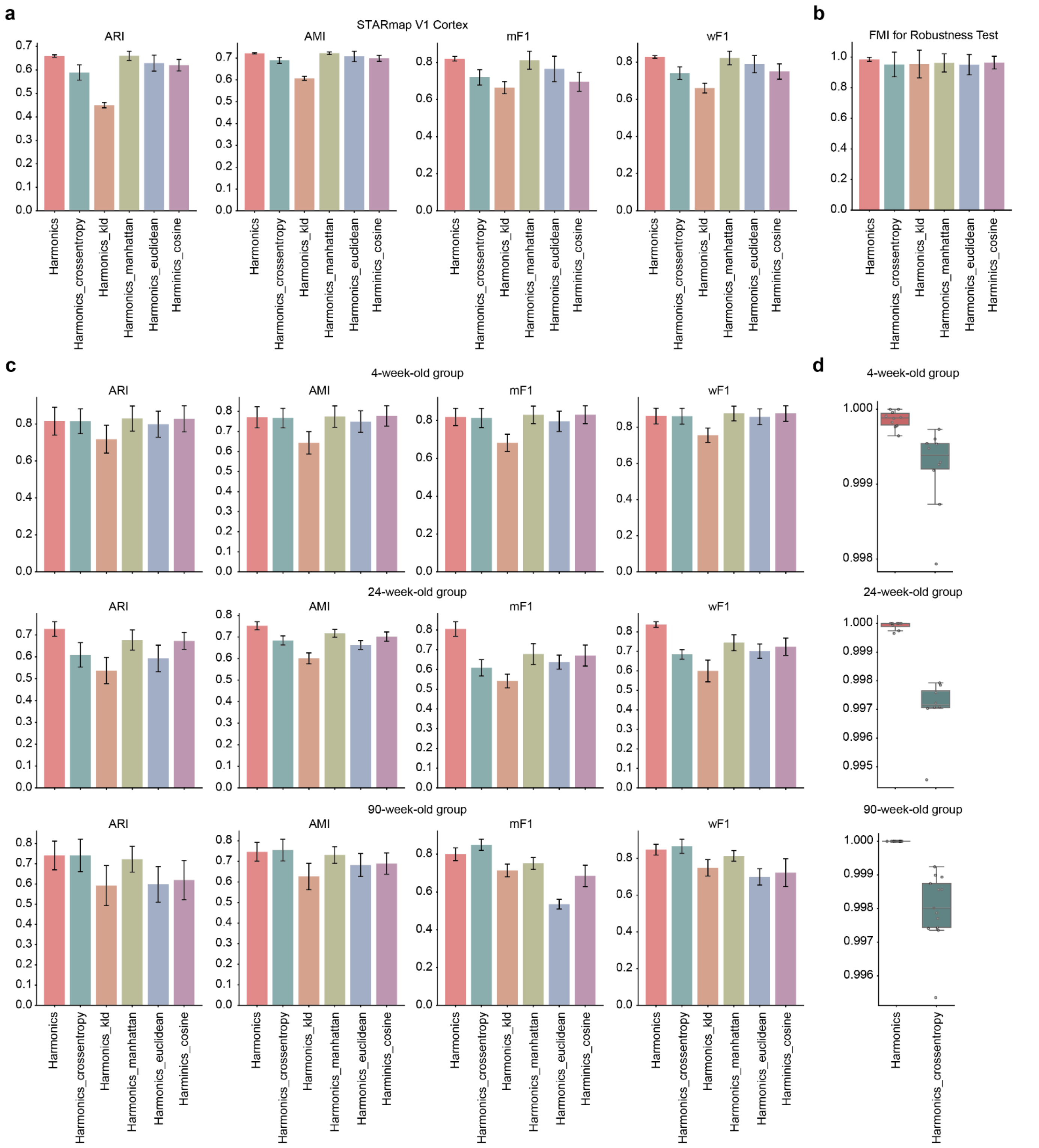

**Supplementary Fig. 2 | Comparison between JSD and commonly used metrics. a, c,** comparison of ARI, AMI, mF1, and wF1 scores for Harmonics using different metrics, including JSD, KLD, cross-entropy (equivalent to the reverse KLD), Manhattan distance, Euclidean distance, and cosine similarity, on the STARmap V1C dataset (**a**) and the MERFISH FC&S dataset (**c**). **b,** robustness test of JSD and other metrics based on pairwise FMI scores across 20 repeated runs on the STARmap V1C dataset. **d,** robustness comparison between JSD and cross-entropy on the MERFISH FC&S dataset, where both metrics show comparable performance in niche identification across two groups of slices. Each dot represents an individual slice. For all box plots, the center line indicates the median; box limits represent the upper and lower quartiles; whiskers extend to the largest and smallest values within 1.5 times the interquartile range. For all bar plots, the height of each bar represents the mean value across repeated experiments (**a**), across all pairwise comparison (**b**), or across all slices (**c**), and the error bars indicate the standard deviation.

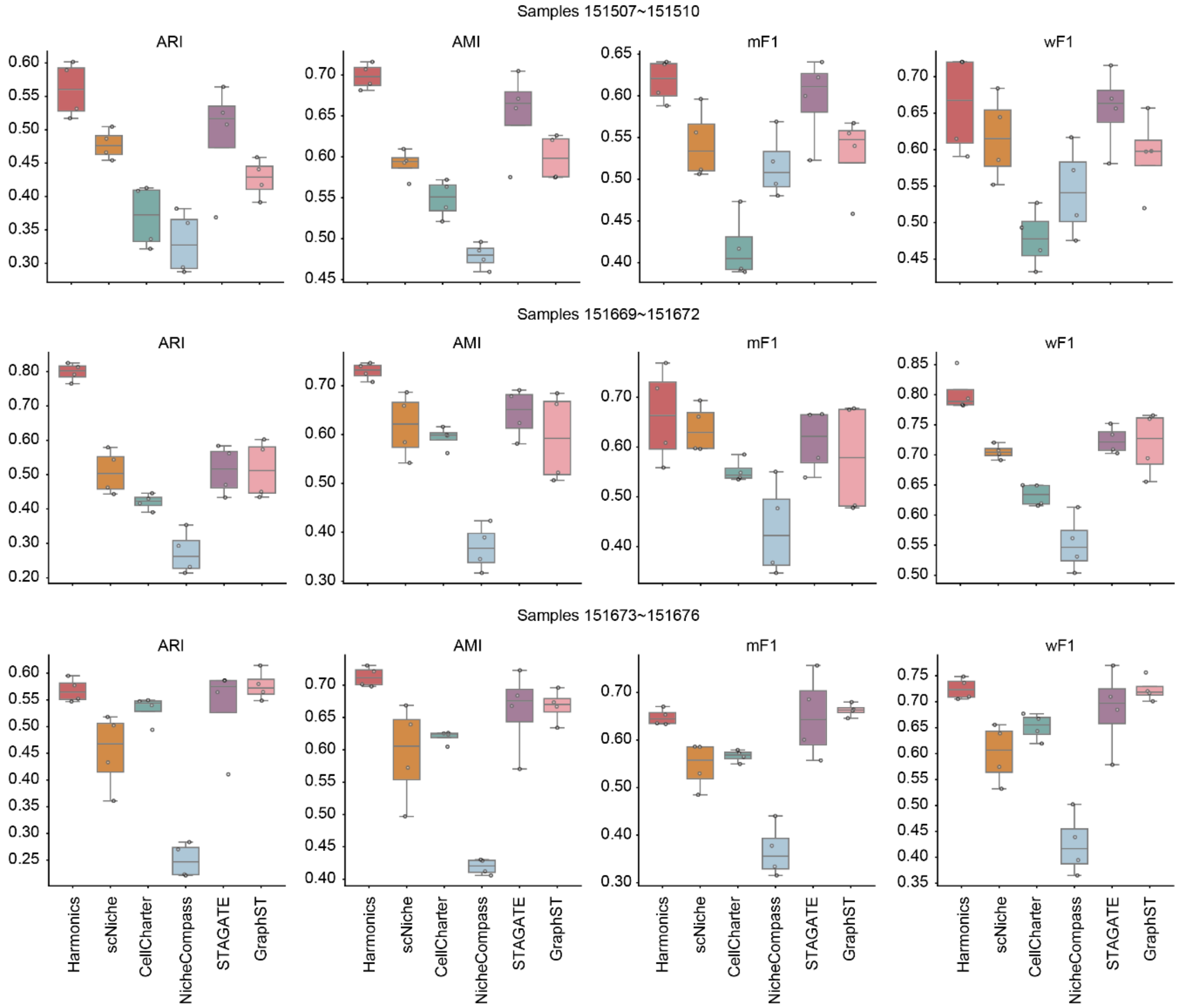

**Supplementary Fig. 3 | Performance evaluation of Harmonics and baseline methods on the Visium DLPFC datasets.** ARI, AMI, mF1, and wF1 are used as evaluation metrics across the three groups. Harmonics achieves superior performance on the first two groups and performs comparably to STAGATE and GraphST on the third. Each dot represents an individual slice. For all box plots, the center line indicates the median; box limits represent the upper and lower quartiles; whiskers extend to the largest and smallest values within 1.5 times the interquartile range.

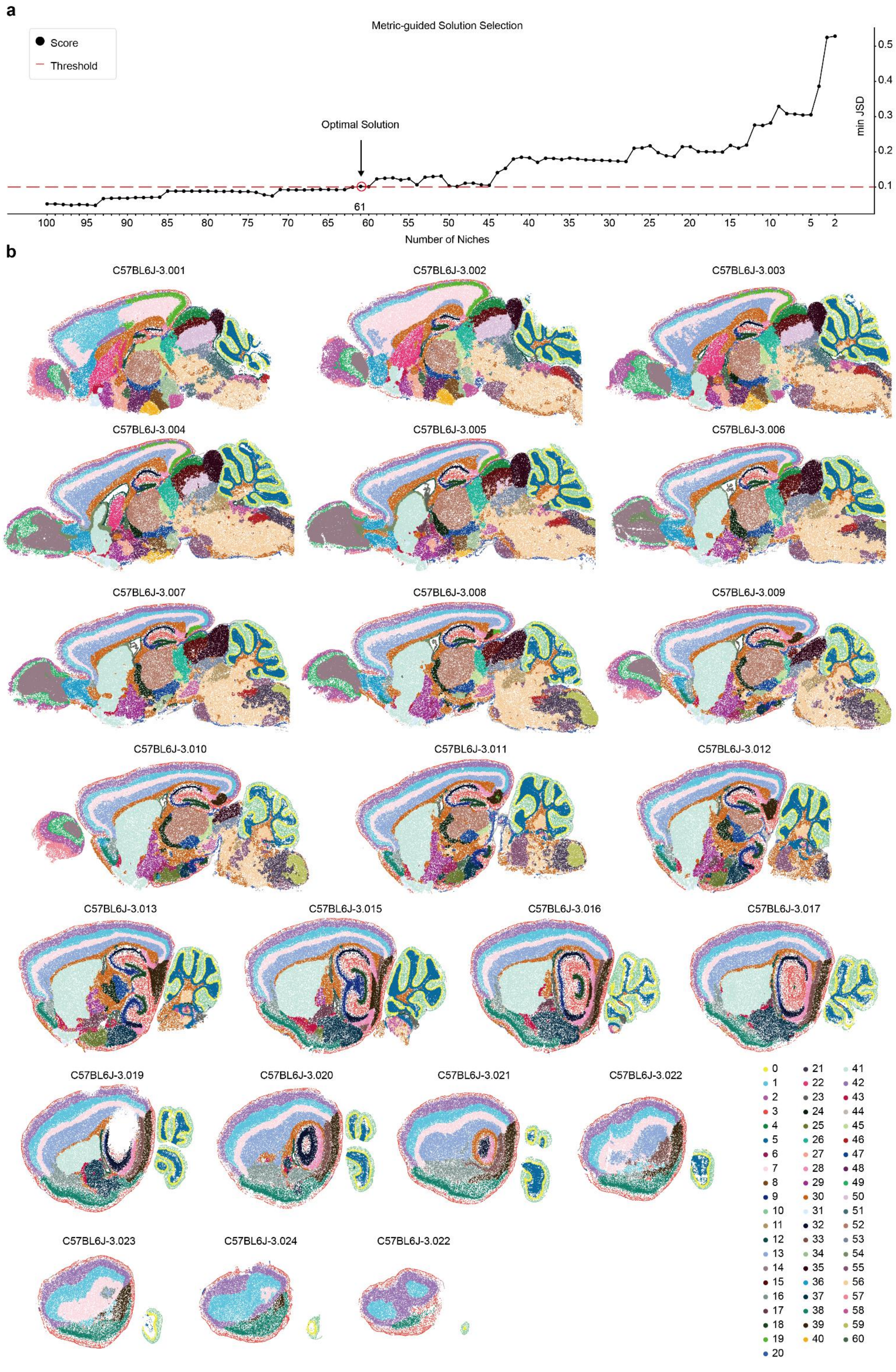
a
Metric-guided Solution Selection
Score
Threshold
Optimal Solution
61
min JSD
0.1
0.2
0.3
0.4
0.5
Number of Niches
100
95
90
85
80
75
70
65
60
55
50
45
40
35
30
25
20
15
10
5
2
b
C57BL6J-3.001
C57BL6J-3.002
C57BL6J-3.003
C57BL6J-3.004
C57BL6J-3.005
C57BL6J-3.006
C57BL6J-3.007
C57BL6J-3.008
C57BL6J-3.009
C57BL6J-3.010
C57BL6J-3.011
C57BL6J-3.012
C57BL6J-3.013
C57BL6J-3.015
C57BL6J-3.016
C57BL6J-3.017
C57BL6J-3.019
C57BL6J-3.020
C57BL6J-3.021
C57BL6J-3.022
C57BL6J-3.023
C57BL6J-3.024
C57BL6J-3.022

**Supplementary Fig. 4 | Outcomes of Harmonics on the MERFISH WMB dataset. a,** $minJSD$ score curve for automatic niche number selection in the MERFISH WMB dataset. **b,** CN assignment results by Harmonics across the 23 consecutive slices, demonstrating consistent niche definition throughout the spatial series.

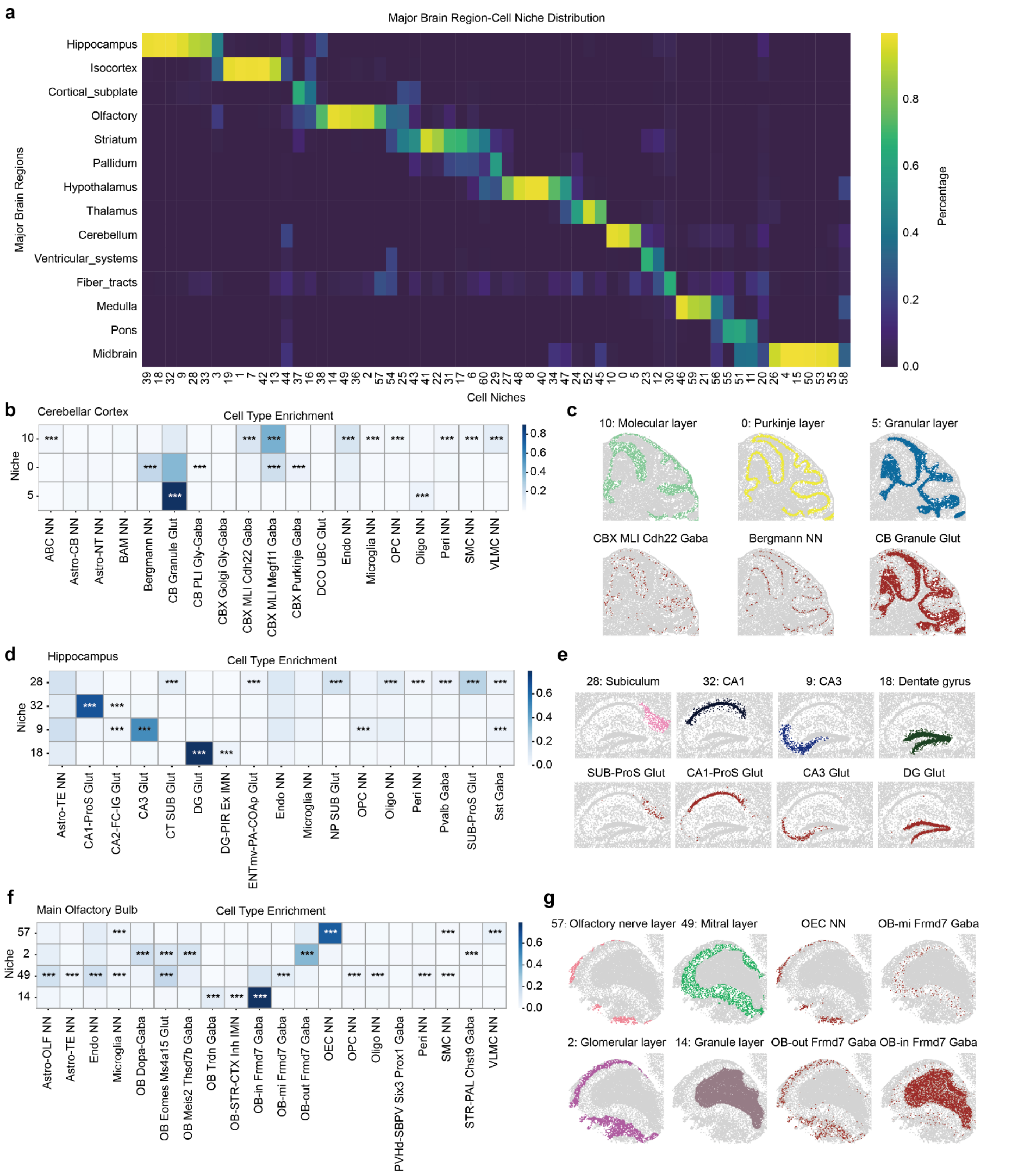

**Supplementary Fig. 5 | Evaluation of Harmonics results on the MERFISH WMB dataset (part 1). a,** distribution of each Harmonics-defined CN across major brain regions, with the unnamed region excluded. Each column is normalized to sum to 1. Most CNs are predominantly localized within a single major region, indicating reasonable CN assignment. **b, d, f,** cell type enrichment analysis for CNs associated with specific brain structures identified by Harmonics, including the cerebellar cortex (CNs 0, 5, and 10) (**b**), hippocampus (CNs 9, 18, 28, and 32) (**d**), and main olfactory bulb (CNs 2, 14, 49, and 57) (**f**). Only cell types with more than 500 cells in the corresponding brain region are included in the analysis. *, adjusted $p$-value < 0.05; **, adjusted $p$-value < 0.01; ***, adjusted $p$-value < 0.001. **c, e, g,** major CNs and their representative enriched cell types in the cerebellar cortex (**c**), hippocampus (**e**), and main olfactory bulb (**g**).

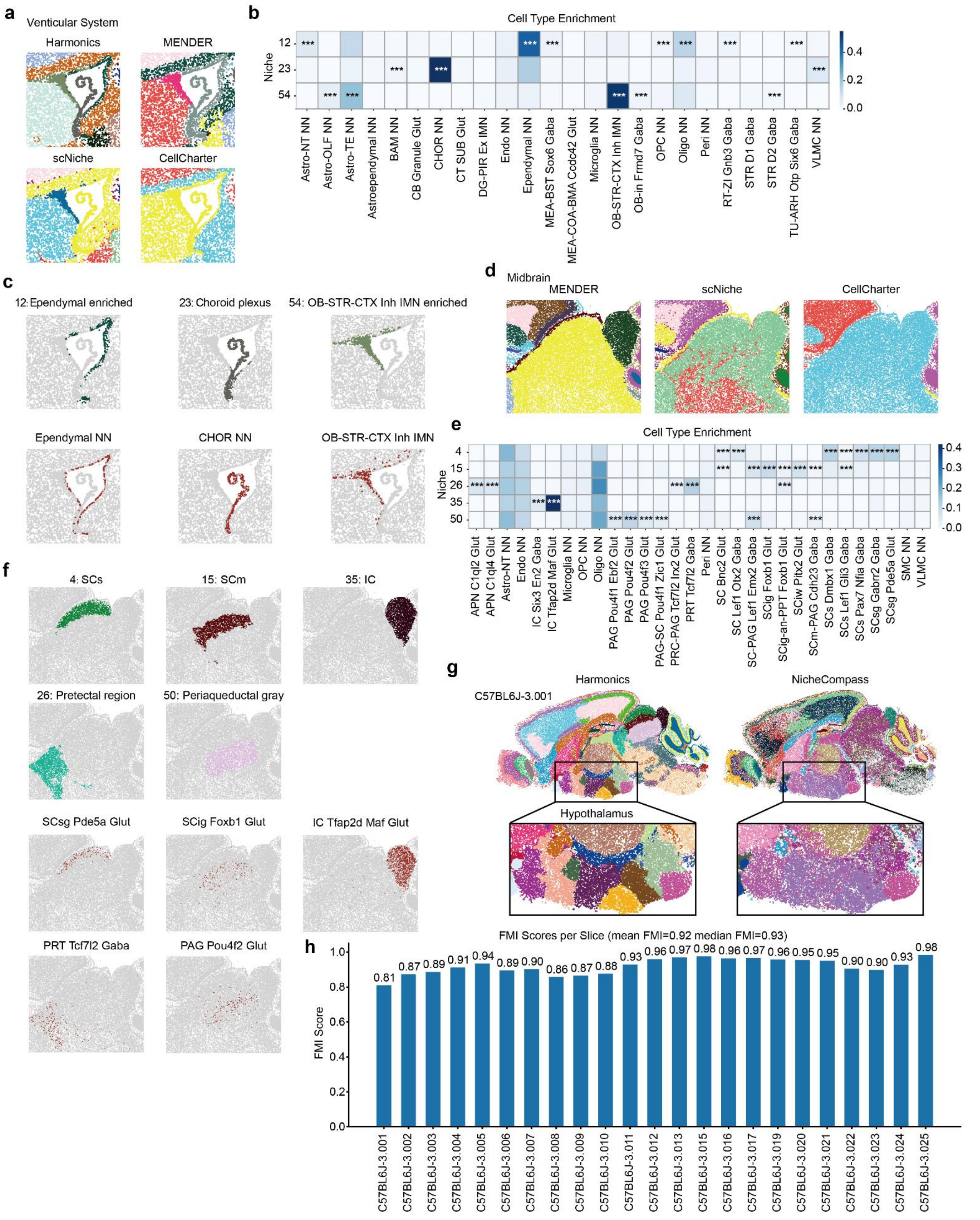

**Supplementary Fig. 6 | Evaluation of Harmonics results on the MERFISH WMB dataset (part 2). a,** comparison of CN identification in the ventricular system, with panels colored by the results of different methods. Harmonics identifies three distinct CNs, whereas baseline methods are unable to distinguish niches at this level of granularity. **b, e,** cell type enrichment analysis for CNs associated with specific brain structures identified by Harmonics, including the ventricular system (CNs 12, 23, and 54) (**b**) and the midbrain (CNs 4, 15, 26, 35, and 50) (**e**). Only cell types with more than 100 cells in the ventricular system (**b**) and more than 500 cells in the midbrain (**e**) are included in the analysis. *, adjusted $p$-value < 0.05; **, adjusted $p$-value <

0.01; ***, adjusted $p$-value < 0.001. **c, f,** major CNs and their representative enriched cell types in the ventricular system (**c**) and midbrain (**f**). SCs, superior colliculus, sensory related; SCm, superior colliculus, motor related; IC, inferior colliculus. **d,** CN definitions produced by baseline methods in the midbrain, with panels colored by the output of different methods. **g,** comparison between Harmonics and NicheCompass in the hypothalamus. Harmonics identifies multiple distinct CNs with clear boundaries, while NicheCompass does not resolve the same level of niche diversity and produces poorly separated regions. **h,** robustness analysis of Harmonics based on label transfer. For each slice, FMI scores are calculated by comparing CN labels in the label transfer setting with those obtained from the full dataset. Both mean and median FMI scores remain high across slices, indicating the strong robustness and consistency of Harmonics.

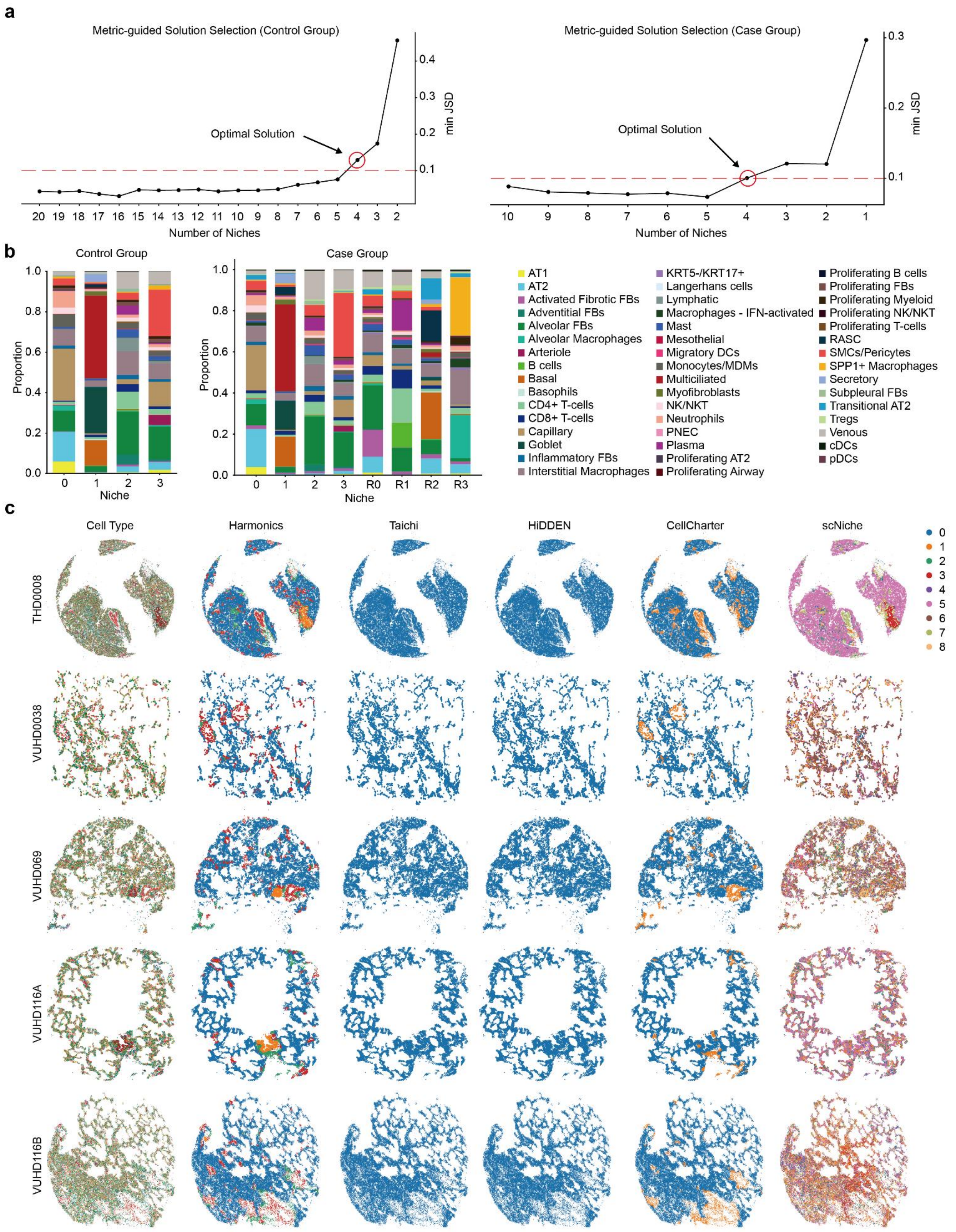

**Supplementary Fig. 7 | Overview of results on the Xenium PF dataset. a,** $minJSD$ score curves used for automatic niche number selection in the control group (left) and case group (right) of the Xenium PF dataset. **b,** cell type compositions across different BCNs in the control group (left) and across different BCNs/CSNs in the case group (right). CSNs defined by Harmonics are labeled with the prefix "R". **c,** representative slices from the control group, colored by cell types as well as niche assignments from Harmonics and baseline methods.

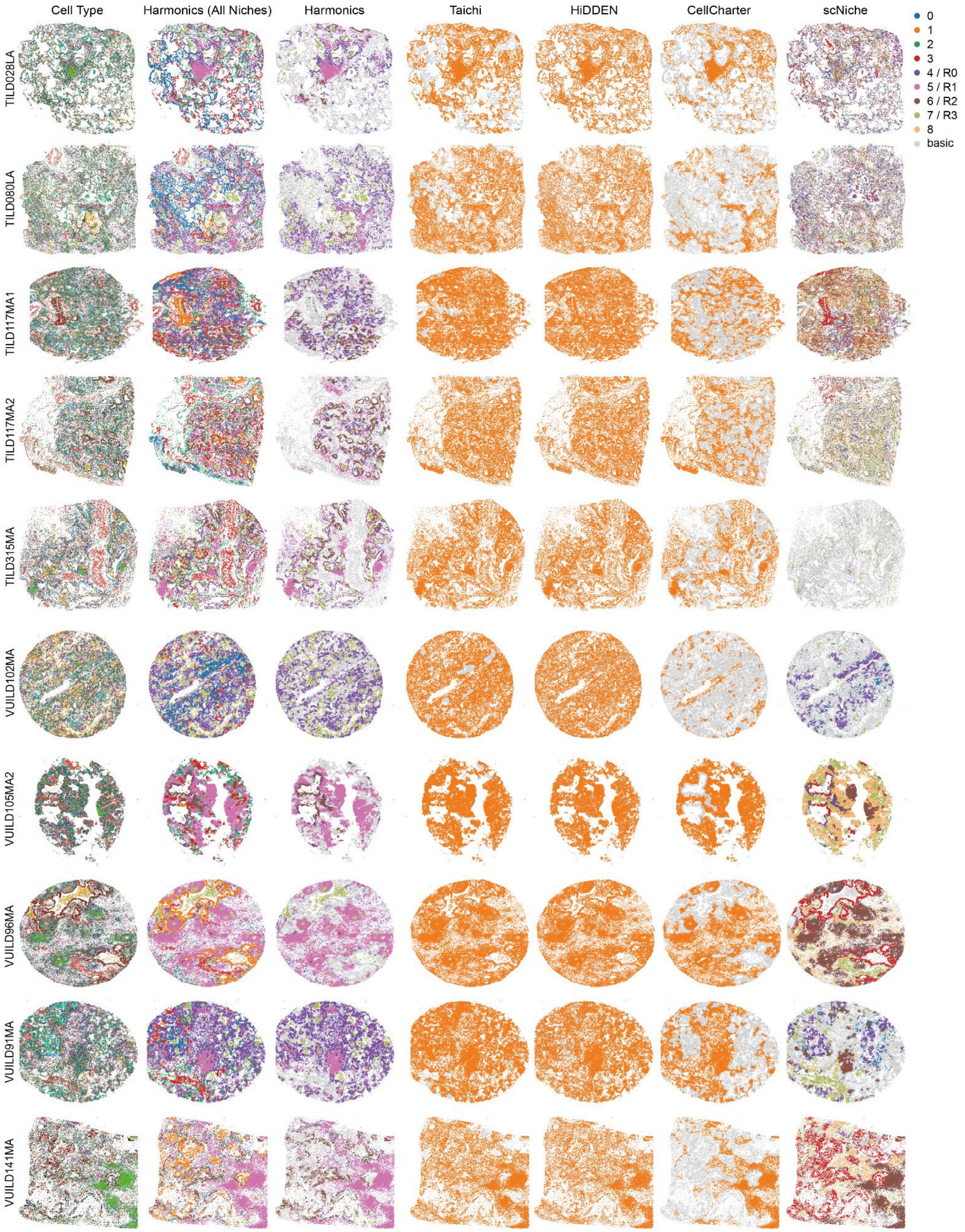

**Supplementary Fig. 8 | Performance of Harmonics and baseline methods in identifying CSNs in the Xenium PF dataset.** Representative slices from the case group of the Xenium PF dataset, shown with cell type annotations, all niches identified by Harmonics, and CSNs defined by Harmonics and baseline methods, where "basic" denotes niches that are not recognized as CSNs. CSNs defined by Harmonics are labeled with the prefix "R".

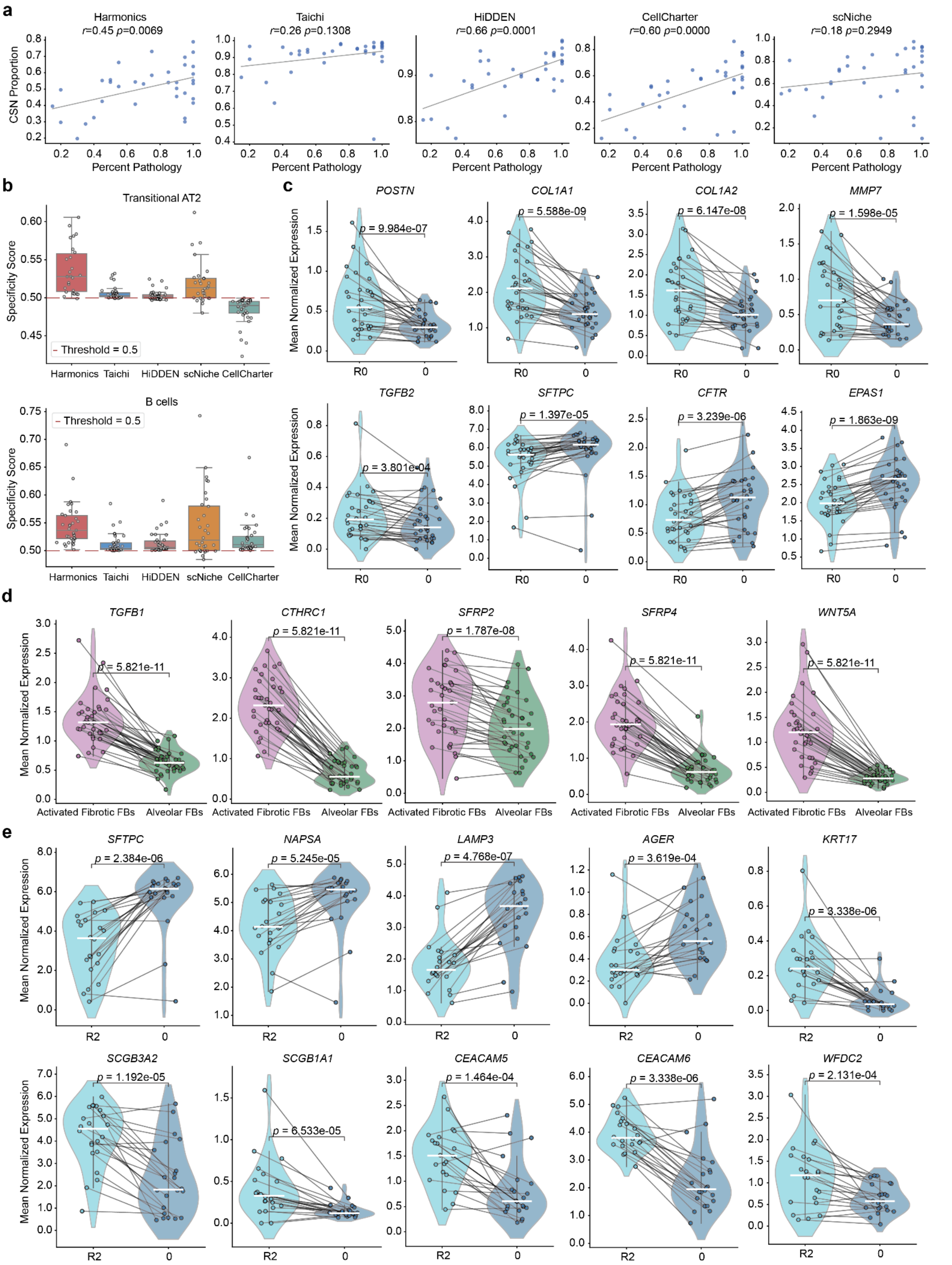

**Supplementary Fig. 9 | Quantitative comparison on the Xenium PF dataset and differential gene expression analysis. a,** correlation between the overall proportion of CSNs identified by each method and pathology percentage. Each dot represents a slice. Trend lines are fitted using univariate linear regression, and Pearson's $r$ along with $p$-values from a two-sided permutation test are reported. Significant associations are observed only for Harmonics, HiDDEN, and CellCharter. **b,** comparison of the specificity of transitional AT2 (top) and B cells (bottom) enrichment in CSNs across all methods. Each dot represents a single slice. For each method, the CSN with the highest median specificity score across all slices is selected for comparison. Dots below the red dashed line represent slices in which the CSN contains a lower proportion of the target cell type than the rest of the tissue, indicating poor specificity. In all box plots, the center line denotes the median; box limits represent the upper and lower quartiles; whiskers extend to the most extreme values within 1.5 times the interquartile range. **c-e,** differential gene expression analyses within the case group: (**c**) AT2 cells between CSN-R0 and BCN-0, (**d**) activated fibrotic FBs versus alveolar FBs within CSN-R0, and (**e**) AT2 cells between CSN-R2 and BCN-0. Each dot represents a slice, and $p$-values are calculated using a two-sided Wilcoxon signed-rank test.

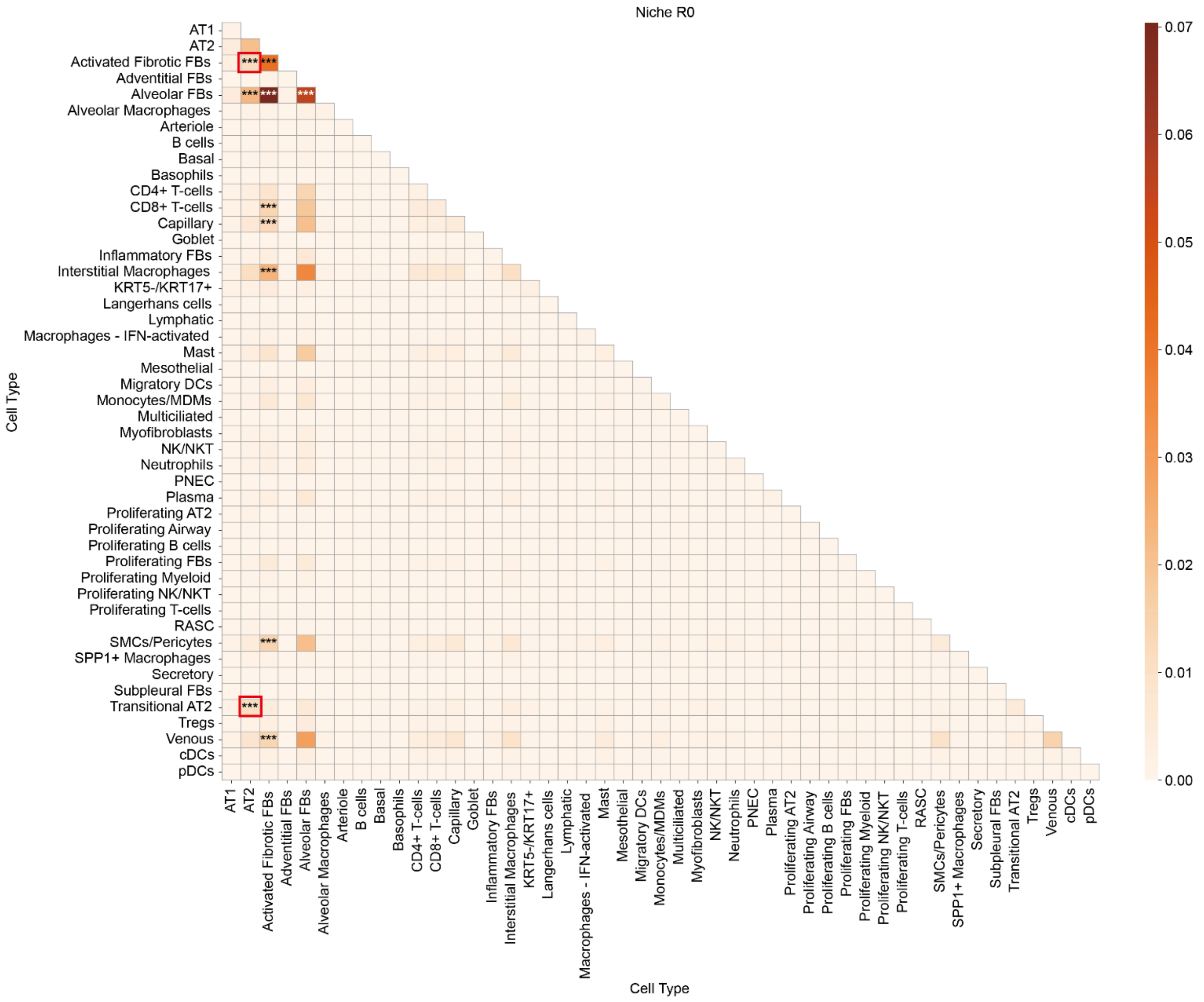

**Supplementary Fig. 10 | Potential CCI enrichment analysis results for CSN-R0 in the Xenium PF dataset.** Spatial interactions between AT2 cells and both activated fibrotic FBs and transitional AT2 cells are significantly enriched in CSN-R0 and highlighted in the figure. *, adjusted $p$-value < 0.05; **, adjusted $p$-value < 0.01; ***, adjusted $p$-value < 0.001.

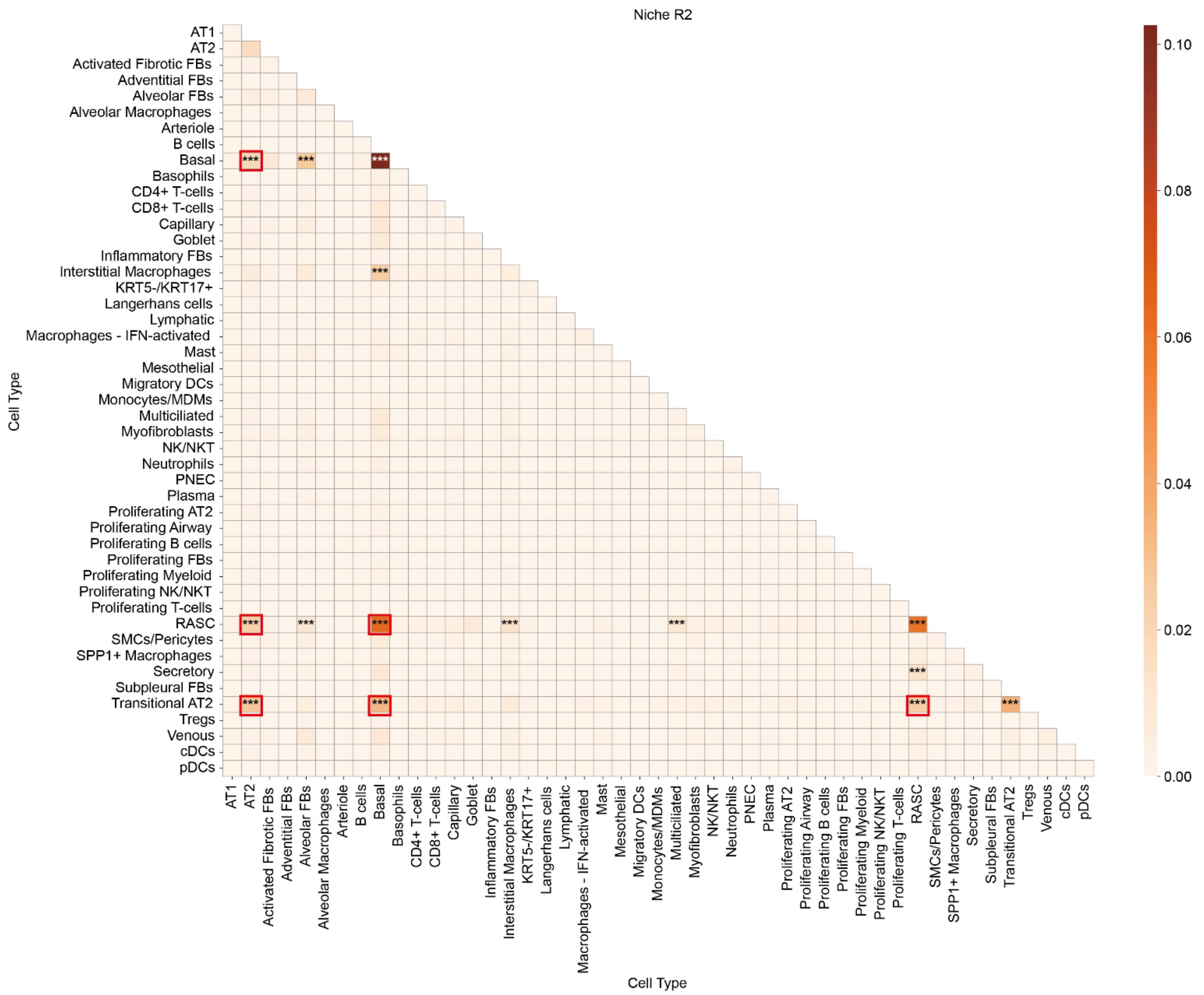

**Supplementary Fig. 11 | Potential CCI enrichment analysis results for CSN-R2 in the Xenium PF dataset.** Pairwise spatial interactions among AT2, basal, RASC, and transitional AT2 cells are significantly enriched in CSN-R2 and highlighted in the figure. *, adjusted $p$-value < 0.05; **, adjusted $p$-value < 0.01; ***, adjusted $p$-value < 0.001.

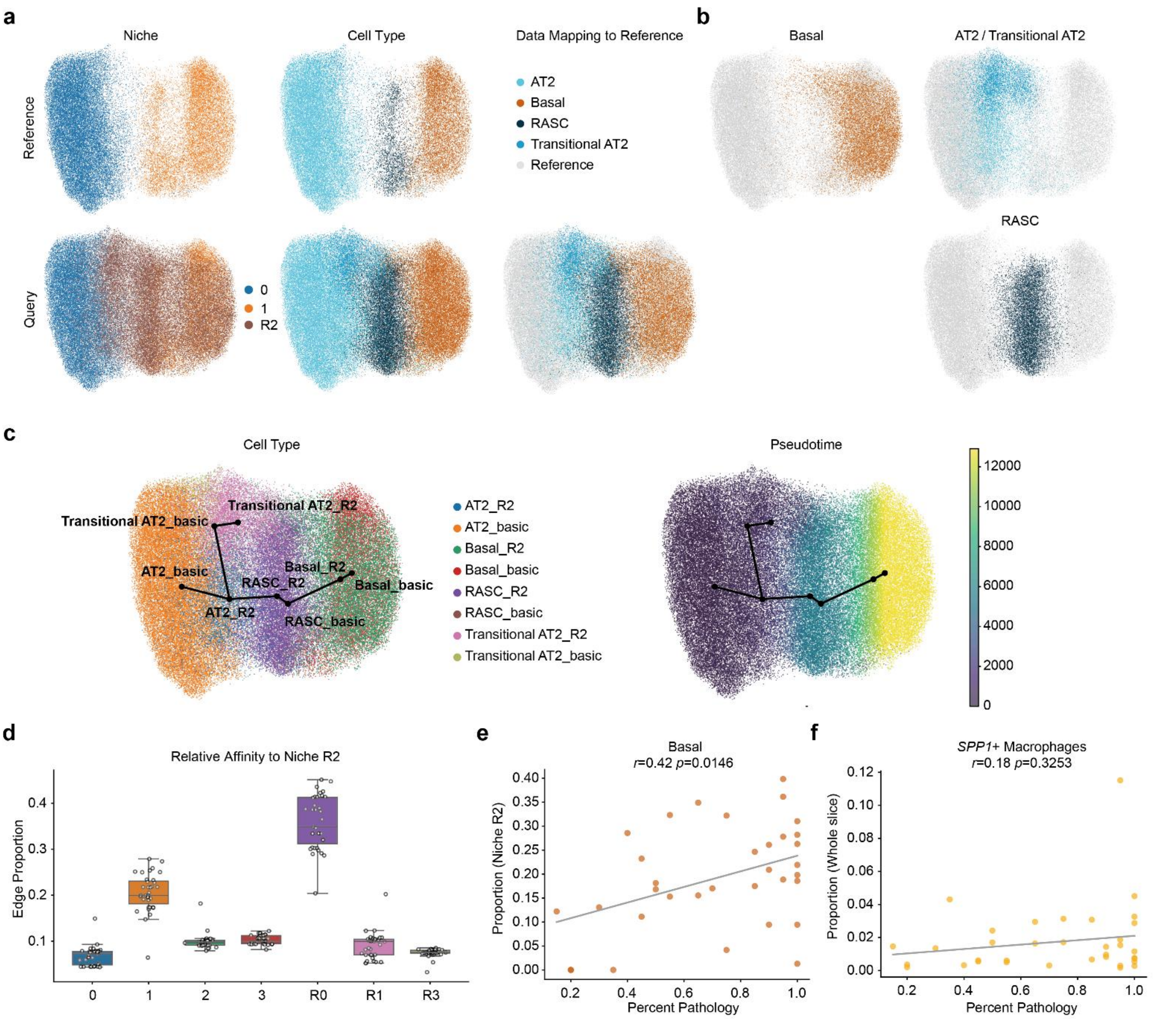

**Supplementary Fig. 12 | Additional results for the analysis of CSN-R2 and CSN-R3 in the Xenium PF dataset. a,** UMAP of AT2, basal, RASC, and transitional AT2 cells from BCN-0 and BCN-1 used as a reference (top), and the projection of the same four cell types from CSN-R2 onto the reference UMAP as query data (right), colored by niche affiliation and cell type. **b,** projection of CSN-R2 cells by cell type, each mapped onto the reference UMAP from (**a**). **c,** trajectory and pseudotime inference of combined reference and query data shown in (**a**), with cells colored by cell type (left) and pseudotime (right). The black lines depict two differentiation trajectories inferred for AT2_basic cells. **d,** comparison of the spatial proximity with CSN-R2. The y-axis indicates the proportion of edges (among all edges connecting CSN-R2 to other niches within the same slice) that are connected to each specific niche. Edges are constructed using Delaunay triangulation and the top 1% longest edges are excluded. Each dot represents a slice. In box plots, the center line denotes the median; box limits represent the upper and lower quartiles; whiskers extend to the most extreme values within 1.5 times the interquartile range. **e, f,** correlation between pathology percentage and the proportion of basal cells within CSN-R2 (**e**) or the overall proportion of *SPP1+* macrophages in the entire slice (**f**). Each dot represents a slice. Trend lines are fitted using univariate linear regression, and Pearson's $r$ along with $p$-values from a two-sided permutation test are reported.

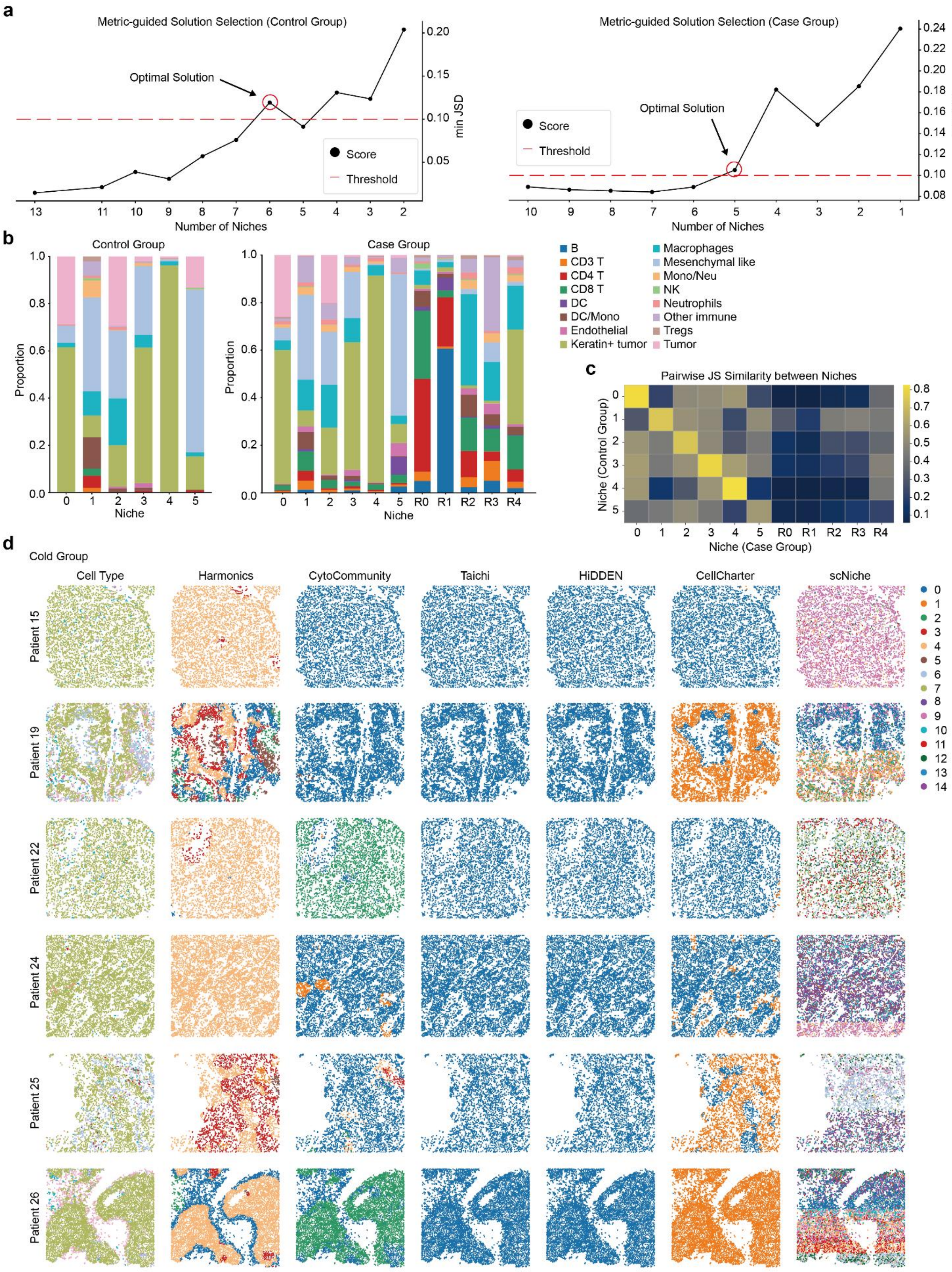

**Supplementary Fig. 13 | Overview of results on the MIBI-TOF TNBC dataset. a,** $minJSD$ score curves used for automatic niche number selection in the control (cold) group (left) and case (compartmentalized and mixed) group (right) of the MIBI-TOF TNBC dataset. **b,** cell type compositions across different niches in the control group (left) and the case group (right). CSNs defined by Harmonics are labeled with the prefix "R". **c,** similarity between each BCN from the control group and each BCN/CSN in the case group, quantified by 1-JSD. **d,** representative slices from the cold group, colored by cell types as well as niche assignments from Harmonics and baseline methods.

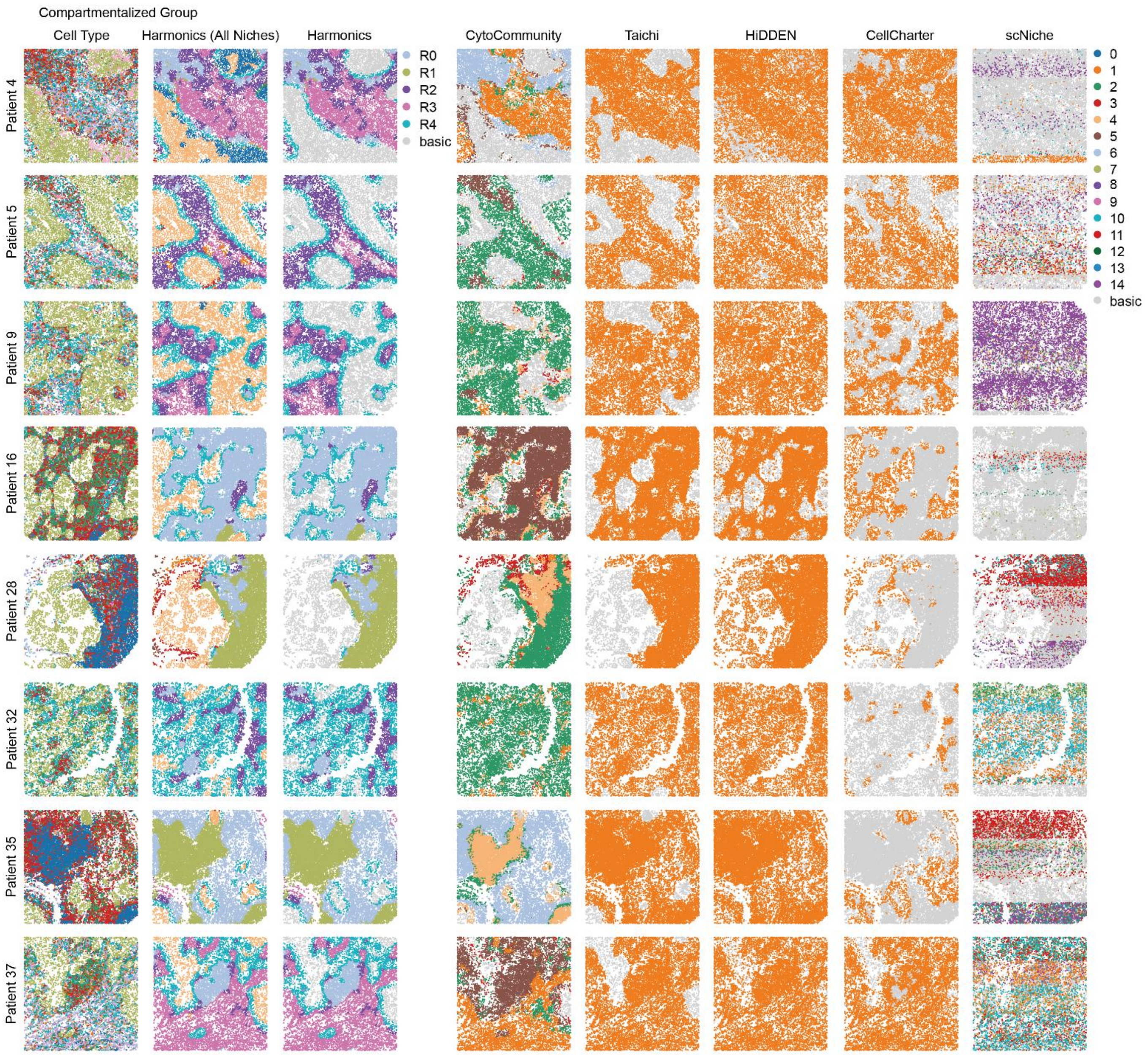

**Supplementary Fig. 14 | Performance of Harmonics and baseline methods in identifying CSNs from the compartmentalized group of the MIBI-TOF TNBC dataset.** Representative slices from the compartmentalized group of the MIBI-TOF TNBC dataset, shown with cell type annotations, all niches identified by Harmonics, and CSNs defined by Harmonics and baseline methods, where "basic" denotes niches that are not recognized as CSNs. CSNs defined by Harmonics are labeled with the prefix "R".

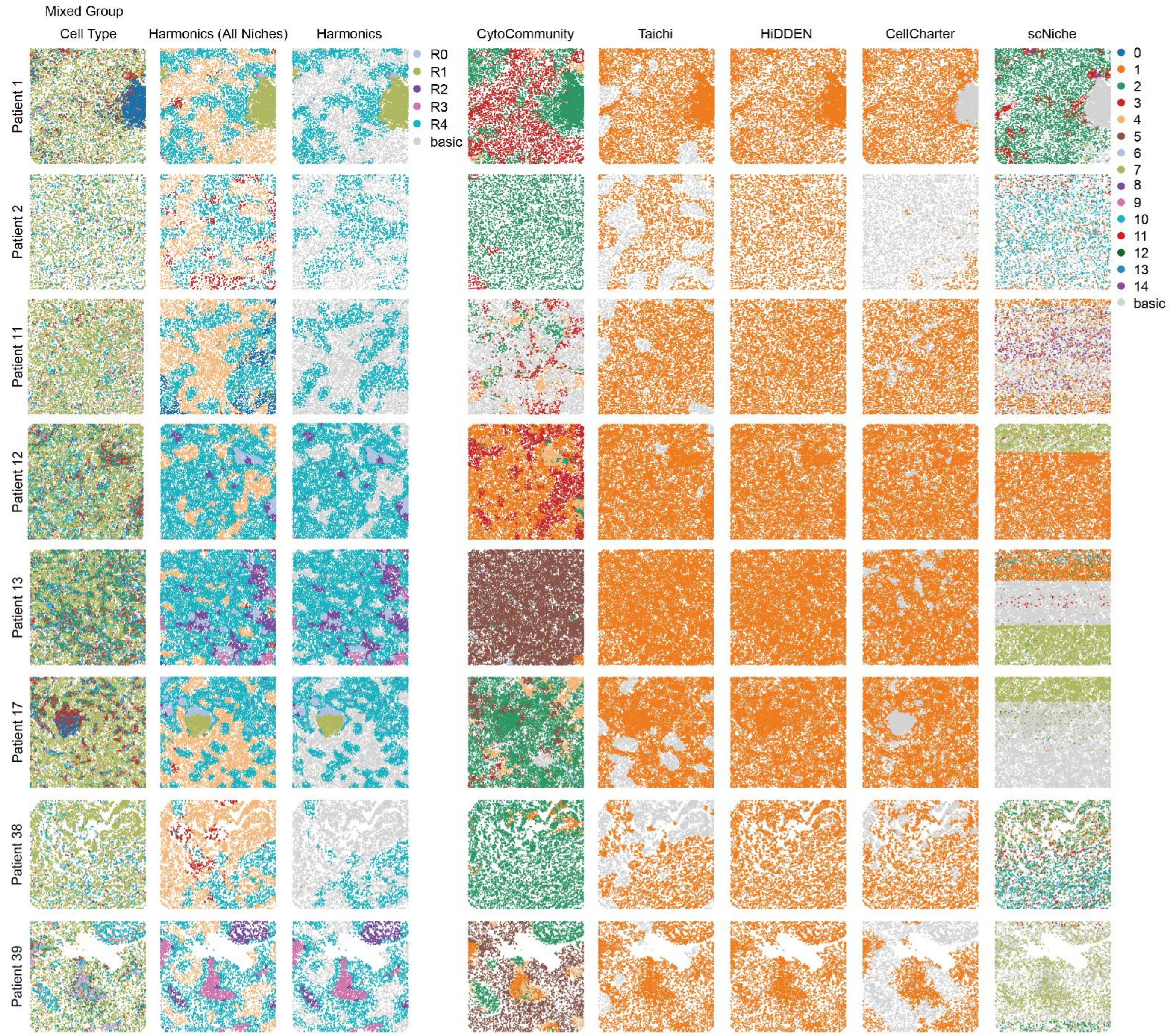

**Supplementary Fig. 15 | Performance of Harmonics and baseline methods in identifying CSNs from the mixed group of the MIBI-TOF TNBC dataset.** Representative slices from the mixed group of the MIBI-TOF TNBC dataset, shown with cell type annotations, all niches identified by Harmonics, and CSNs defined by Harmonics and baseline methods, where "basic" denotes niches that are not recognized as CSNs. CSNs defined by Harmonics are labeled with the prefix "R".

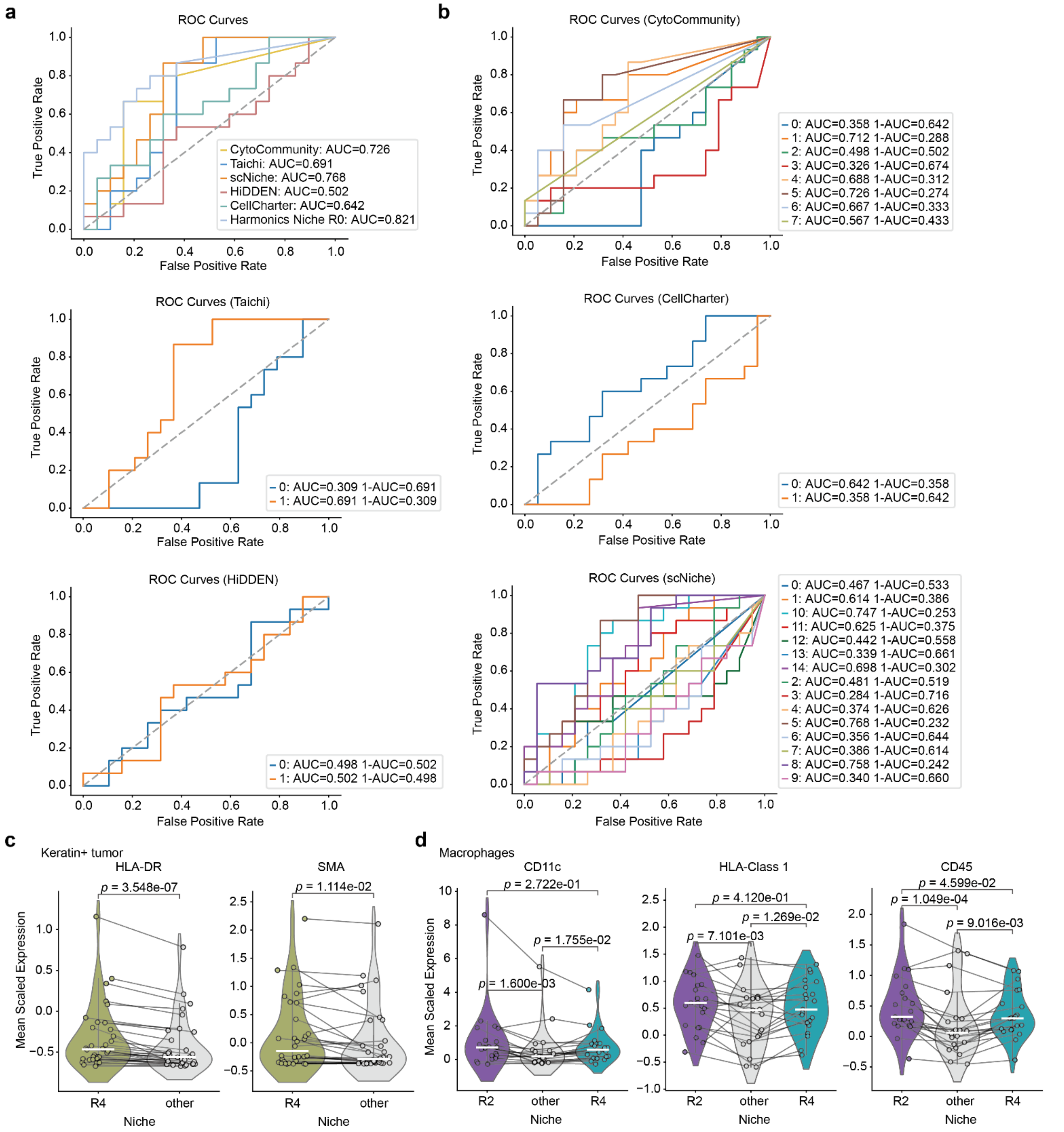

**Supplementary Fig. 16 | Comparison between Harmonics and baseline methods in disease subtype classification. a,** comparison of different methods in distinguishing compartmentalized and mixed groups using niche proportion. The AUC is reported. For Harmonics, the proportion of CSN-R0 is used to classify disease subtypes; for other methods, the niche that can best distinguish between subtypes is selected. **b,** evaluation of each baseline method's ability to distinguish disease subtypes using niche proportions. For each niche, both AUC and 1-AUC are reported. **c,** differential expression of HLA-DR and SMA in Keratin+ tumor cells between CSN-R4 and other niches. **d,** differential expression of CD11c, HLA-Class 1, and CD45 in macrophages between CSN-R2/CSN-R4 and other niches. In panels **c** and **d**, each dot represents a slice, and $p$-values are computed using a two-sided Wilcoxon signed-rank test.

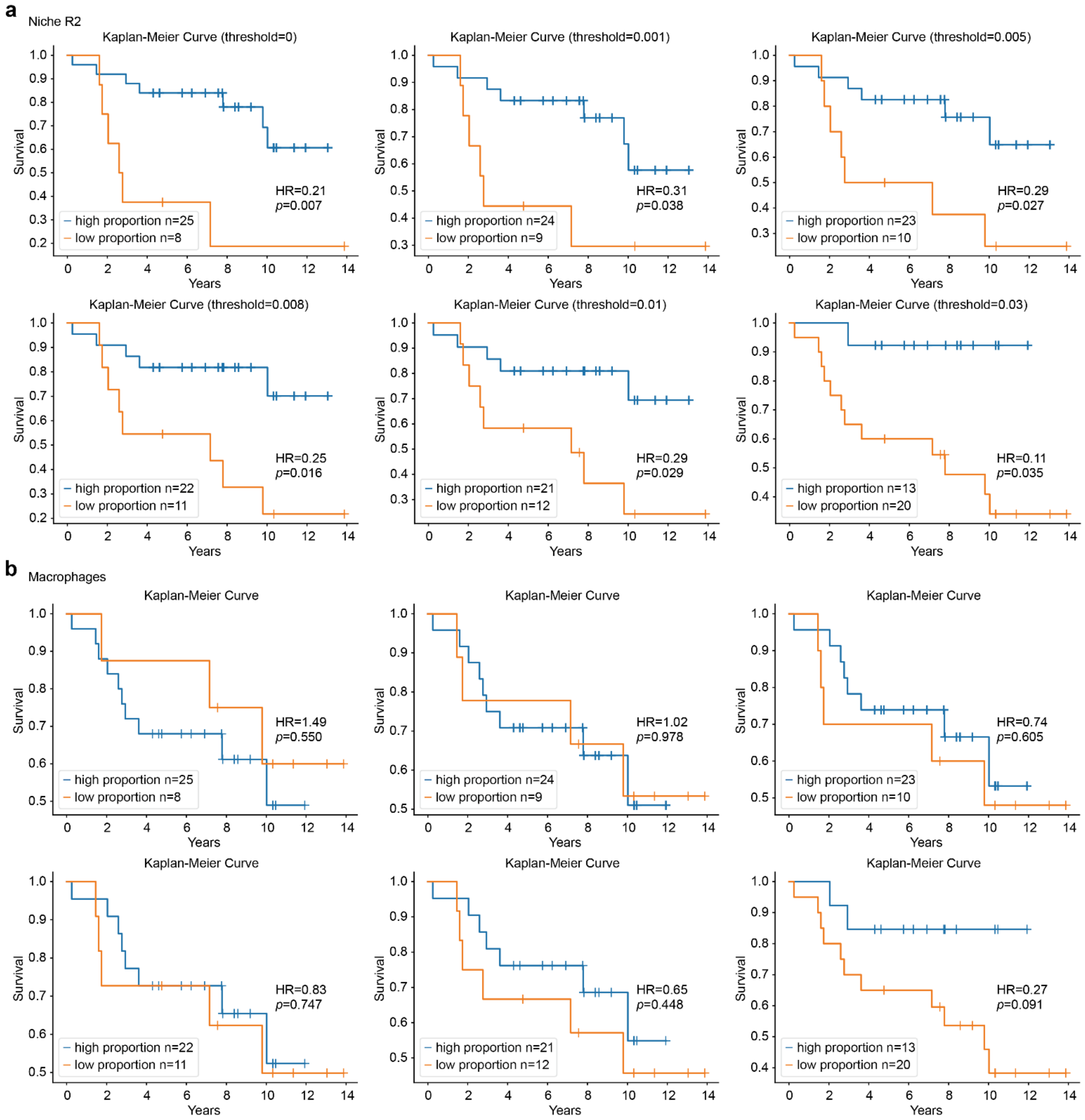

**Supplementary Fig. 17 | Survival analysis using niche-level and cell-level information. a, b,** Kaplan–Meier survival curves for patients stratified by high and low intra-slice proportions of CSN-R2 (**a**) or macrophages, the dominant cell type within CSN-R2 (**b**). The HRs are estimated using Cox proportional hazards regression, and the associated $p$-values are derived from a two-sided test. Stratification thresholds in (**a**) are set to 0, 0.001, 0.005, 0.008, 0.01, and 0.03. In (**b**), thresholds are selected to produce high and low groups with the same patient counts as the corresponding stratifications in (**a**), ensuring comparability of group sizes.

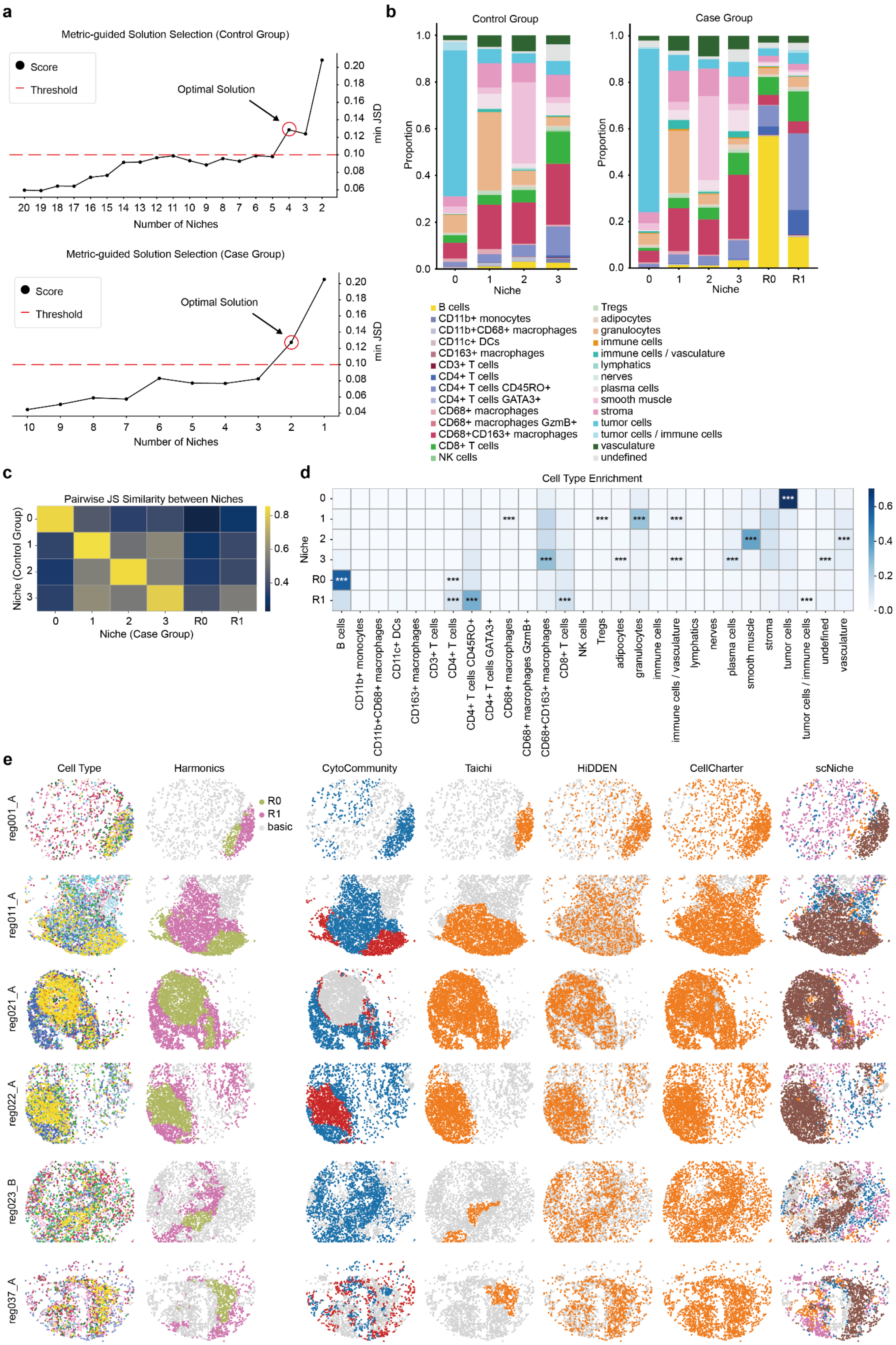
a
Metric-guided Solution Selection (Control Group)
Score
Threshold
Optimal Solution
min JSD
Number of Niches
Metric-guided Solution Selection (Case Group)
b
Control Group
Case Group
Proportion
Niche
B cells
CD11b+ monocytes
CD11b+CD68+ macrophages
CD11c+ DCs
CD163+ macrophages
CD3+ T cells
CD4+ T cells
CD4+ T cells CD45RO+
CD4+ T cells GATA3+
CD68+ macrophages
CD68+ macrophages GzmB+
CD68+CD163+ macrophages
CD8+ T cells
NK cells
Tregs
adipocytes
granulocytes
immune cells
immune cells / vasculature
lymphatics
nerves
plasma cells
smooth muscle
stroma
tumor cells
tumor cells / immune cells
vasculature
undefined
c
Pairwise JS Similarity between Niches
Niche (Control Group)
Niche (Case Group)
d
Cell Type Enrichment
Niche
e
Cell Type
Harmonics
CytoCommunity
Taichi
HiDDEN
CellCharter
scNiche
R0
R1
basic
reg001_A
reg011_A
reg021_A
reg022_A
reg023_B
reg037_A

**Supplementary Fig. 18 | Performance evaluation on the CODEX CRC dataset. a,** $minJSD$ score curves used for automatic determination of niche numbers in the control (DII) group (top) and the case (CLR) group (bottom). **b,** cell type compositions of different niches in the control (left) and case (right) groups. CSNs identified by Harmonics are labeled with the prefix "R". **c,** similarity between each BCN from the control group and each BCN/CSN in the case group, quantified by 1-JSD. **d,** cell type enrichment analysis for the case group. *, adjusted $p$-value < 0.05; **, adjusted $p$-value < 0.01; ***, adjusted $p$-value < 0.001. **e,** representative slices from the CLR group of the CODEX CRC dataset, shown with cell type annotations, as well as CSNs defined by Harmonics and baseline methods, where "basic" denotes niches that are not recognized as CSNs.

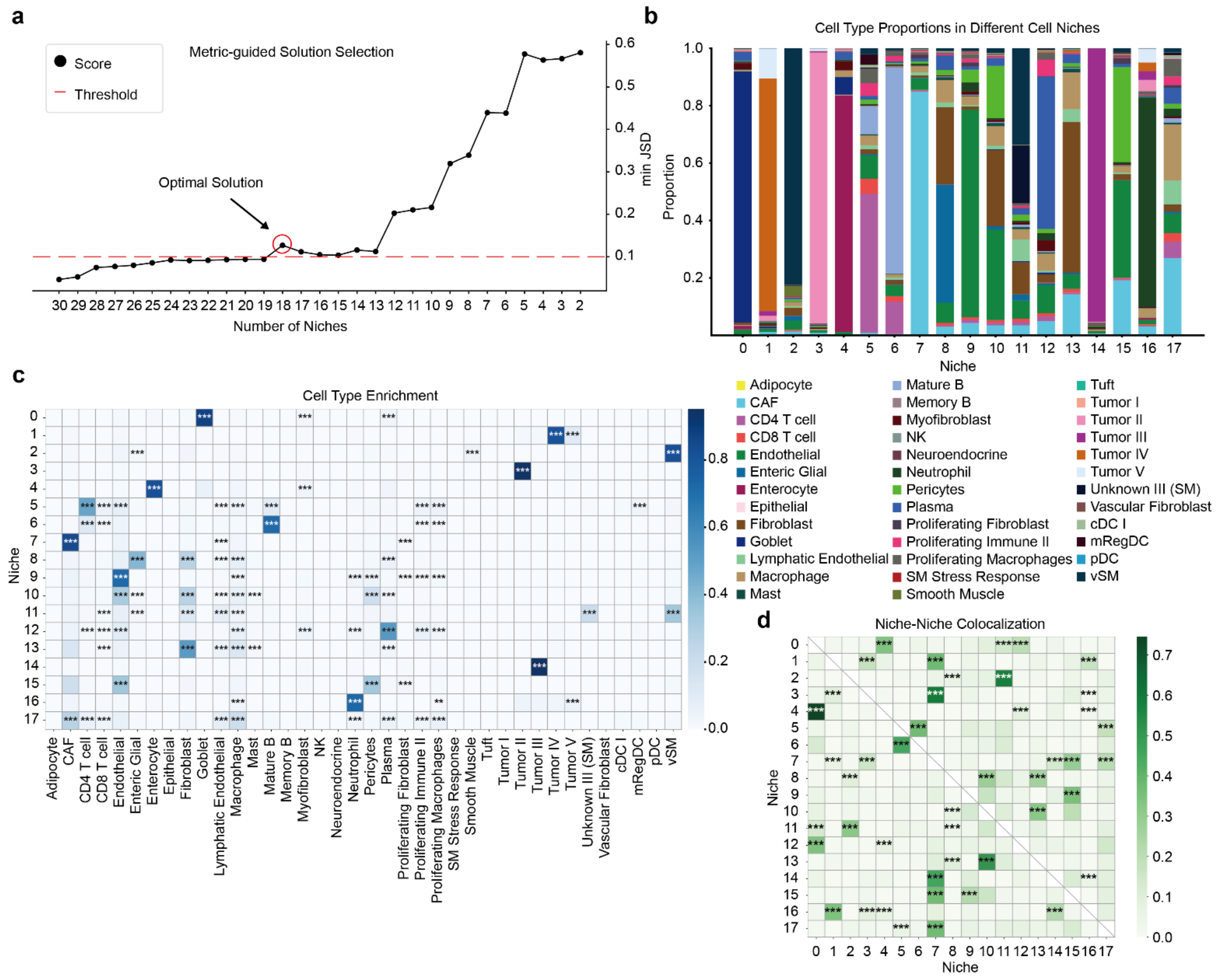

**Supplementary Fig. 19 | Overall results of Harmonics on the Visium HD CRC dataset. a,** $minJSD$ score curve used for automatic determination of niche numbers. **b,** cell type compositions of different niches. **c,** cell type enrichment analysis results. **d,** NNC patterns identified by Harmonics in the Visium HD CRC dataset. Rows represent source niches and columns represent target niches. A significant value indicates that the target niche is significantly enriched in the spatial neighborhood of the source niche. *, adjusted $p$-value < 0.05; **, adjusted $p$-value < 0.01; ***, adjusted $p$-value < 0.001.

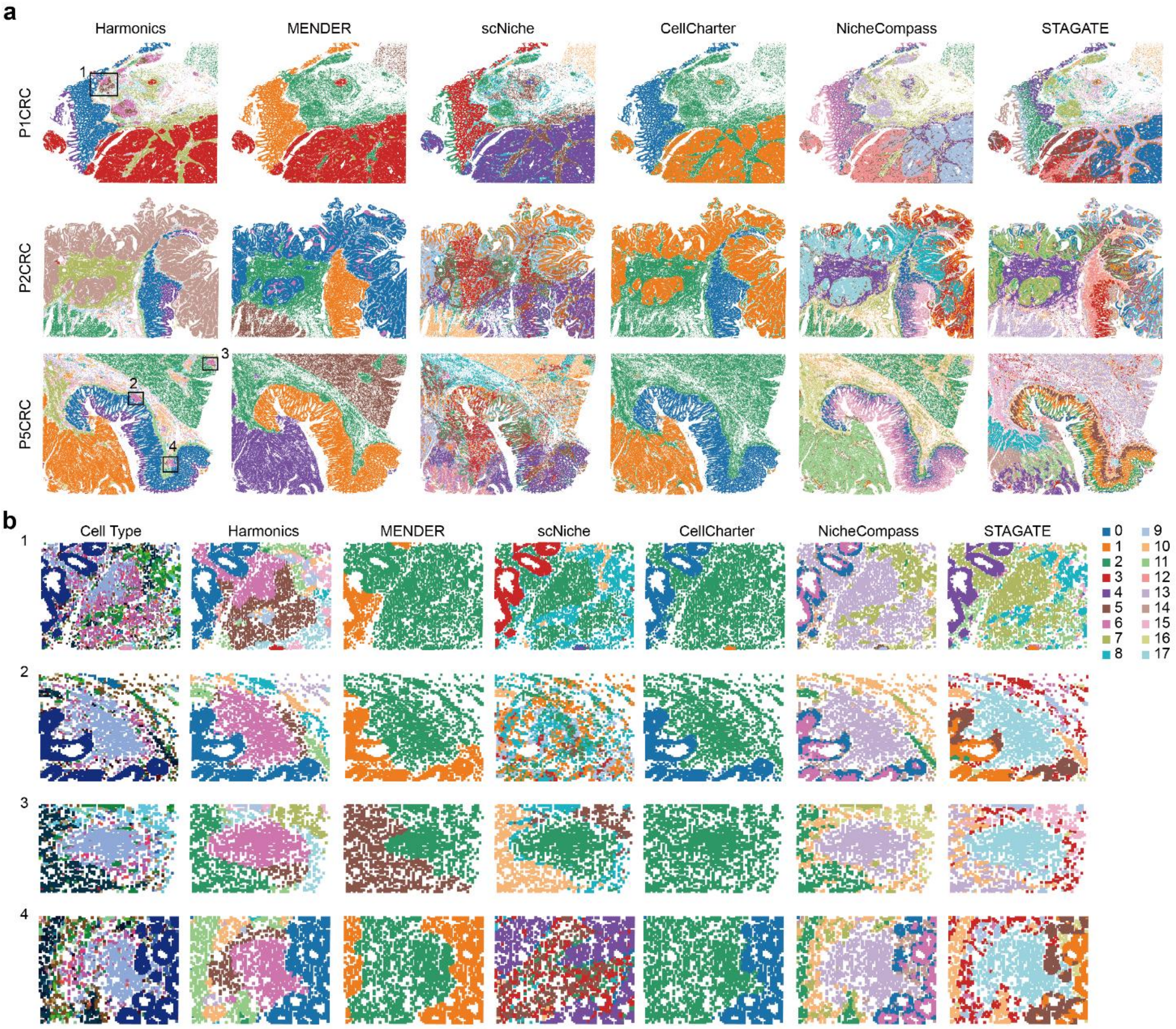

**Supplementary Fig. 20 | Comparison of Harmonics and baseline methods on the Visium HD CRC dataset. a,** CN assignments derived from Harmonics and baseline methods. Four representative ROIs containing TLSs are outlined. **b,** comparison of CN assignments within the four ROIs. Only Harmonics successfully distinguishes the B cell zone from the T cell zone.

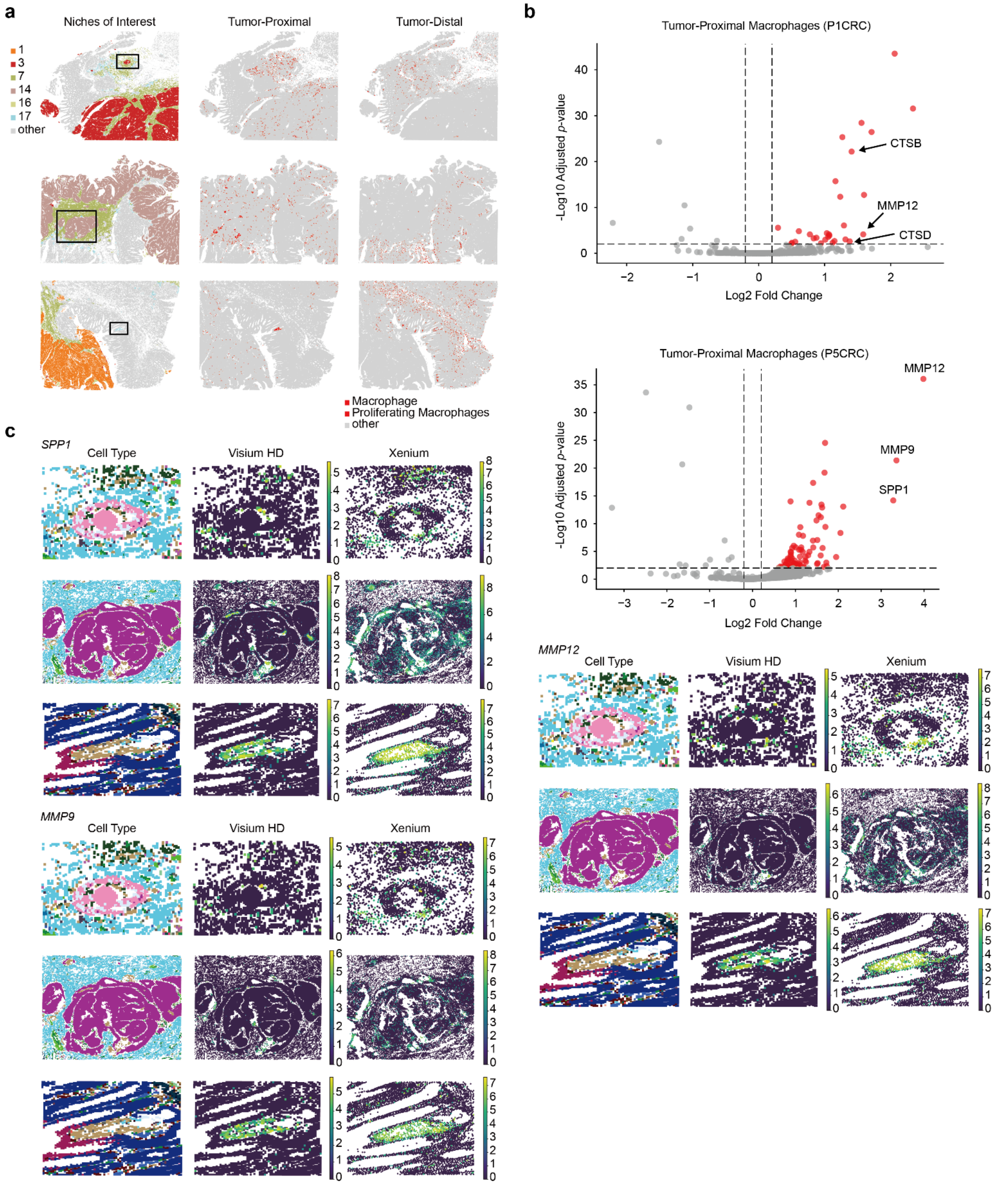

**Supplementary Fig. 21 | Differential analysis of tumor-proximal and tumor-distal macrophages. a,** spatial distribution of macrophages in three representative slices. One ROI is outlined for each slice. The slices are colored by CNs of interest (left), tumor-proximal macrophages (middle), and tumor-distal macrophages (right). **b,** volcano plots showing differentially expressed genes in tumor-proximal macrophages from the P1CRC and P5CRC slices. Genes with an adjusted *p*-values below 0.01 and log2 fold change above 0.2 are considered significantly upregulated and highlighted in red. **c,** expression patterns of three representative upregulated genes, namely *SPP1*, *MMP9*, and *MMP12*, in tumor-proximal macrophages across the ROIs. Both Visium HD and corresponding Xenium sections are shown.
